# CTAT-LR-fusion: accurate fusion transcript identification from long and short read isoform sequencing at bulk or single cell resolution

**DOI:** 10.1101/2024.02.24.581862

**Authors:** Qian Qin, Victoria Popic, Houlin Yu, Emily White, Akanksha Khorgade, Asa Shin, Kirsty Wienand, Arthur Dondi, Niko Beerenwinkel, Francisca Vazquez, Aziz M. Al’Khafaji, Brian J. Haas

## Abstract

Gene fusions are found as cancer drivers in diverse adult and pediatric cancers. Accurate detection of fusion transcripts is essential in cancer clinical diagnostics, prognostics, and for guiding therapeutic development. Most currently available methods for fusion transcript detection are compatible with Illumina RNA-seq involving highly accurate short read sequences. Recent advances in long read isoform sequencing enable the detection of fusion transcripts at unprecedented resolution in bulk and single cell samples. Here we developed a new computational tool CTAT-LR-fusion to detect fusion transcripts from long read RNA-seq with or without companion short reads, with applications to bulk or single cell transcriptomes. We demonstrate that CTAT-LR-fusion exceeds fusion detection accuracy of alternative methods as benchmarked with simulated and real long read RNA-seq. Using short and long read RNA-seq, we further apply CTAT-LR-fusion to bulk transcriptomes of nine tumor cell lines, and to tumor single cells derived from a melanoma sample and three metastatic high grade serous ovarian carcinoma samples. In both bulk and in single cell RNA-seq, long isoform reads yielded higher sensitivity for fusion detection than short reads with notable exceptions. By combining short and long reads in CTAT-LR-fusion, we are able to further maximize detection of fusion splicing isoforms and fusion-expressing tumor cells. CTAT-LR-fusion is available at https://github.com/TrinityCTAT/CTAT-LR-fusion/wiki.

## Introduction

Genomic rearrangements involving chromosomal translocations or deletions can yield fusion genes, in some cases activating oncogenes or disabling tumor suppressors and contributing to cancer. While most cancer relevant fusion genes are found at low levels of recurrence in surveys of diverse tumor types, certain fusions represent hallmark drivers of cancer found at high levels of recurrence, such as BCR::ABL1 in chronic myelogenous leukemia (CML) (Kurzrock et al. 1988), SS18::SSX (Ren et al. 2013) in synovial sarcoma, and TMPRSS2::ERG (Wang et al. 2017) in prostate cancer. Several gene fusions serve as diagnostic markers for certain pediatric cancers, including EWSR1::FLI1 for Ewing’s sarcoma (May et al. 1993), ETV6::RUNX1 in acute lymphoblastic leukemia (Sundaresh and Williams 2017), and PVT1::MYC in medulloblastoma (Northcott et al. 2012), PAX3::FOXO1 in rhabdomyosarcoma (Linardic 2008). The molecular mechanisms by which gene fusions contribute to cancer can widely vary from positioning the 3’ fused gene under the promoter and gene expression regulatory elements of the 5’ gene, or encoding fusion proteins with altered molecular functions, all leading to alterations in the cellular circuitry that ultimately drive uncontrolled cellular proliferation.

Identification of the gene fusions has been an essential part of charting the landscape of cancer genomic variations, deriving biomarkers for molecular diagnostics of cancer patients, and targeting therapies such as tyrosine kinase inhibitors for the treatment of kinase gene fusions such as BCR::ABL1 in CML patients (Cuellar et al. 2018) and EML4::ALK (Christopoulos et al. 2018) in lung cancer. Transcribed and translated gene fusions are of particular interest towards discovering neoantigens in targeted immunotherapies (Yang et al. 2019), yielding additional opportunities for targeting immunotherapies towards cancers with low mutational burdens.

During the past decade, RNA-seq has been the preferred assay for comprehensive gene fusion detection due to its lower cost than whole genome sequencing (WGS) and directly measuring the transcripts arising from the gene fusions. Illumina short-read RNA-seq has become routine for such studies, and numerous computational methods have been developed to predict fusions from Illumina RNA-seq (Kim and Salzberg 2011; Li et al. 2011; McPherson et al. 2011; Benelli et al. 2012; Jia et al. 2013; Wang et al. 2013; Davidson et al. 2015; Latysheva and Babu 2016; Okonechnikov et al. 2016; Rodriguez-Martin et al. 2017; Akers et al. 2018; Haas et al. 2019; Uhrig et al. 2021). Primarily through studies of lllumina RNA-seq, large catalogs of fusions have been cataloged across large collections of tumor and normal tissues (Klijn et al. 2015; Yoshihara et al. 2015; Babiceanu et al. 2016; Hu et al. 2018; Dehghannasiri et al. 2019; Haas et al. 2023). Fusion transcripts relevant to cancer tend to involve genome rearrangements, whereas fusion transcripts identified in normal tissues tend to derive from cis-or trans-splicing or otherwise derive from natural population structural variants yielding population-specific cis-spliced fusion transcripts (Nigro et al. 1991; Li et al. 2008; Li et al. 2009; Chase et al. 2010; Boettger et al. 2012; Qin et al. 2015).

While short RNA-seq reads have been highly useful for identifying fusion gene candidates and resolving fusion transcript isoform breakpoints, the reads are not long enough to resolve the complete isoforms that are expressed, and additional transcript reconstruction methods are needed to infer potential full-length fusion transcripts. Short read RNA-seq methods that involve targeted sequencing of the 3’ or 5’ terminus of RNA molecules, which are currently standard in high throughput single cell sequencing assays, pose further limitations for fusion detection as short reads are less likely to cover the breakpoint of the fusion transcript.

Long read isoform sequencing is made possible by PacBio and Oxford Nanopore Technologies (ONT), enabling full-length isoform sequences via their cDNA, or in the case of ONT, the option of direct RNA sequencing. Early applications of these technologies have been constrained due to low throughput and high error rates. Recent advances in both long-read platforms have enabled high throughput long read transcriptome sequencing at high sequencing accuracy (on par or exceeding that of conventional short read sequencing) (Wenger et al. 2019; Marx 2023). Applications of long isoform reads have enabled deeper insights into transcriptome isoform diversity in whole tissues (Glinos et al. 2022; Reese et al. 2023), and most recently for single cells (Al’Khafaji et al. 2023). Applications of long read RNA-seq is gaining traction in the cancer research community, particularly involving fusion isoform detection, with several computational methods now available that are specifically tailored towards characteristics of long reads (Liu et al. 2020; Davidson et al. 2022; Chen et al. 2023). However, as long read isoform sequencing technology has been rapidly advancing and most computational tools for fusion detection have only recently been developed, there has been limited work thus far towards benchmarking their capabilities or applying them in new areas such as fusion detection in single cells.

To further advance fusion transcript detection using long read isoform sequencing, we developed a new method as part of the Trinity Cancer Transcriptome Analysis Toolkit (CTAT) called CTAT-LR-fusion. CTAT-LR-fusion is specifically developed for long read RNA-seq with or without short read RNA-seq as a modularized software that contains chimeric read extraction, fusion transcripts identification, expression quantification, gene fusion annotation and interactive visualization. To benchmark existing tools, we collected or generated comprehensive simulation datasets to reflect varied sequencing technologies and sequencing error rates. We also designed new experiments to profile a normal cell line transcriptome with spiked-in known oncogenic fusion transcripts and nine cancer cell lines using the same long read sequencing protocol MAS-ISO-seq (Al’Khafaji et al. 2023). In both simulation and real datasets, we systematically benchmarked CTAT-LR-fusion accuracy in comparison to available long read fusion tools, demonstrating top performance in each setting. We finally applied CTAT-LR-fusion to long isoform read sequences derived from tumor single cell transcriptomes including melanoma and high grade serous ovarian carcinoma (HGSOC) metastases, in each case discovering fusion transcripts that distinguished tumor and normal cell states. In all experiments with real data, we used available sample-matched Illumina short reads or generated companion Illumina RNA-seq for comparison to long isoform reads and to augment findings based on long reads. CTAT-LR-fusion is freely available as an open-source software at https://github.com/TrinityCTAT/CTAT-LR-fusion/wiki.

## Results

### CTAT-LR-fusion pipeline

Fusion transcript detection from long reads by CTAT-LR-fusion involves two phases (**Figure 1a**). In the first phase, candidate chimeric long reads are rapidly identified using a customized version of the minimap2 aligner (Li 2018) that only reports alignments for reads with preliminary mappings to multiple genomic loci. Candidate chimeric reads and corresponding fusion gene pairs are identified based on these preliminary alignments. In the second phase, candidate fusion gene pairs are modeled as collinear gene contigs by FusionInspector (Haas et al. 2023) (included with CTAT-LR-fusion), and the candidate chimeric reads are realigned to the fusion contigs using minimap2 full alignment. Fusion genes are identified based on high quality read alignments and fusion transcript breakpoints quantified based on the number of supporting long isoform fusion reads (see **Methods** for details). If sample-matched Illumina RNA-seq is available, FusionInspector is further executed to capture short read alignment evidence for these fusion candidates, and the FusionInspector results are integrated with the long read results into the final CTAT-LR-Fusion report. Long reads (and with short reads where applicable) alignment evidence for fusion transcripts is made available for further navigation via the included interactive web-based IGV-report (**Figure 1b**) or separately via desktop IGV (Robinson et al. 2011).

**Figure 1:**
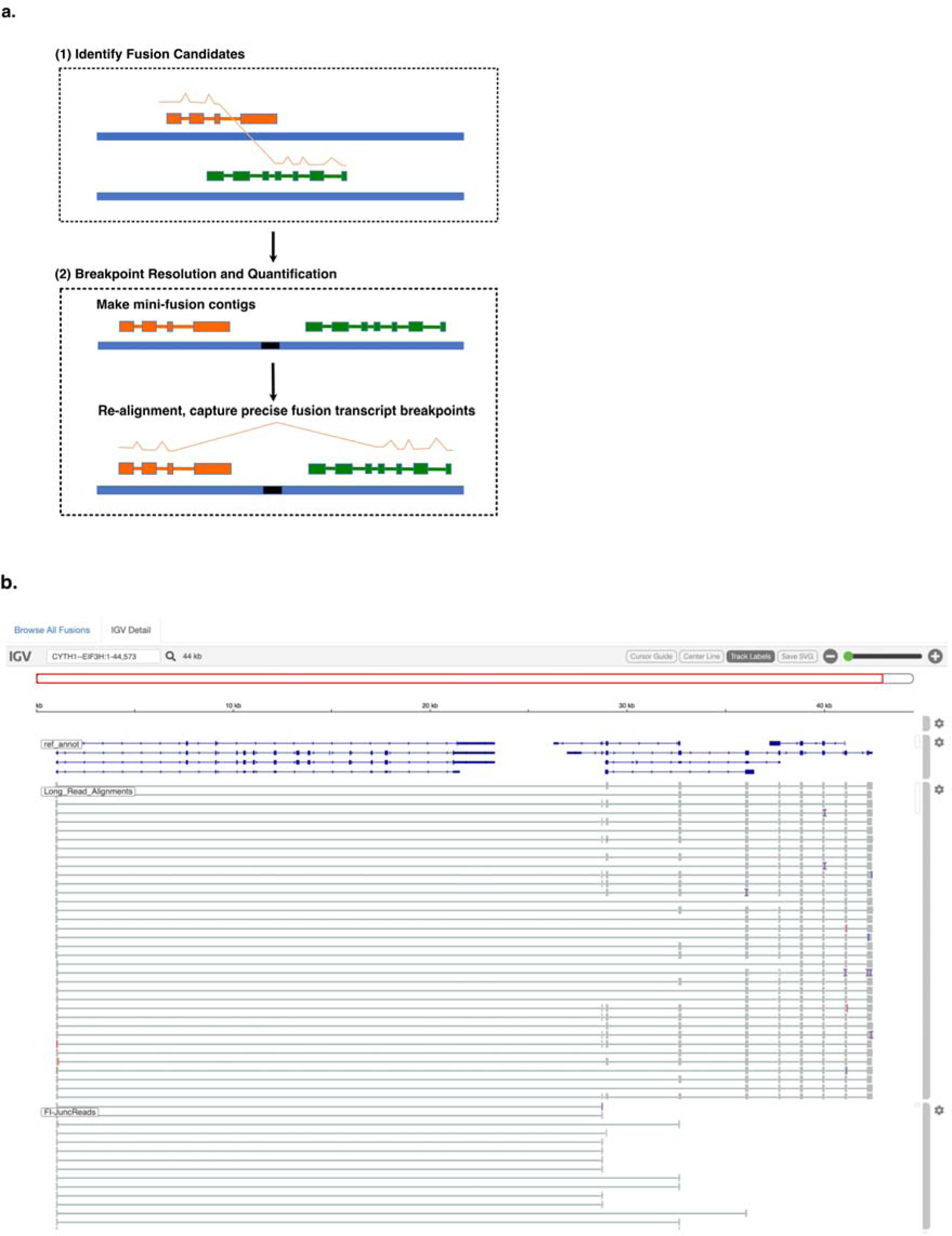
CTAT-LR-fusion algorithm and output. (a) CTAT-LR-fusion workflow. (b) IGV-reports visualization providing interactive analysis of long isoform read alignment evidence for predicted fusion transcripts, including alignments for matched Illumina short reads where available.

### Fusion Transcript Detection Accuracy Using Simulated Long Reads

Earlier benchmarking of fusion transcript detection by JAFFAL (Davidson et al. 2022) entailed the use of BadRead (Wick 2019) to simulate long reads for fusion transcripts based on PacBio and ONT error models and spanning a wide range of sequence divergence from 25% error (75% alignment identity) to 5% error (95% alignment identity). We leveraged these available test data to examine CTAT-LR-fusion accuracy in comparison to available alternatives, including JAFFAL (Davidson et al. 2022), LongGF (Liu et al. 2020), FusionSeeker (Chen et al. 2023), and pbfusion (Roger Volden 2023).

For each long read fusion transcript detection method, we computed precision, recall, and corresponding F1 accuracy score according to minimum read support, and captured the maximum accuracy for each test data set representative of sequencing technology (PacBio or ONT) and error rate (75% to 95% sequence identity) (**Figure 2a,b**). Surprisingly, only CTAT-LR-fusion, JAFFAL, and pbfusion (since version 0.4.0) properly report fusion gene pairs in the order in which they are fused together from 5’ to 3’ in the corresponding fusion transcript, and so only CTAT-LR-fusion, JAFFAL, and pbfusion exhibit high accuracy when benchmarking fusion detection in a ‘strict’ manner requiring ordered gene pairs. Relaxing this requirement and scoring fusion detection based solely on unordered gene pairings, all methods demonstrate moderate to high fusion detection accuracy at the lowest sequence divergence (95% identity) for both PacBio and ONT simulated reads. Unsurprisingly, fusion detection accuracy improves with read sequence quality for all methods. In comparison to the other methods, pbfusion was most sensitive to error rates and least capable of fusion detection with the highest error rates and largely incompatible with the divergent ONT simulated reads. Overall, CTAT-LR-fusion and JAFFAL were found to be top-performing with these simulated test data when considering fusion gene order and orientation, with CTAT-LR-fusion providing top-performance across most combinations of error rates and sequencing technology.

**Figure 2.**
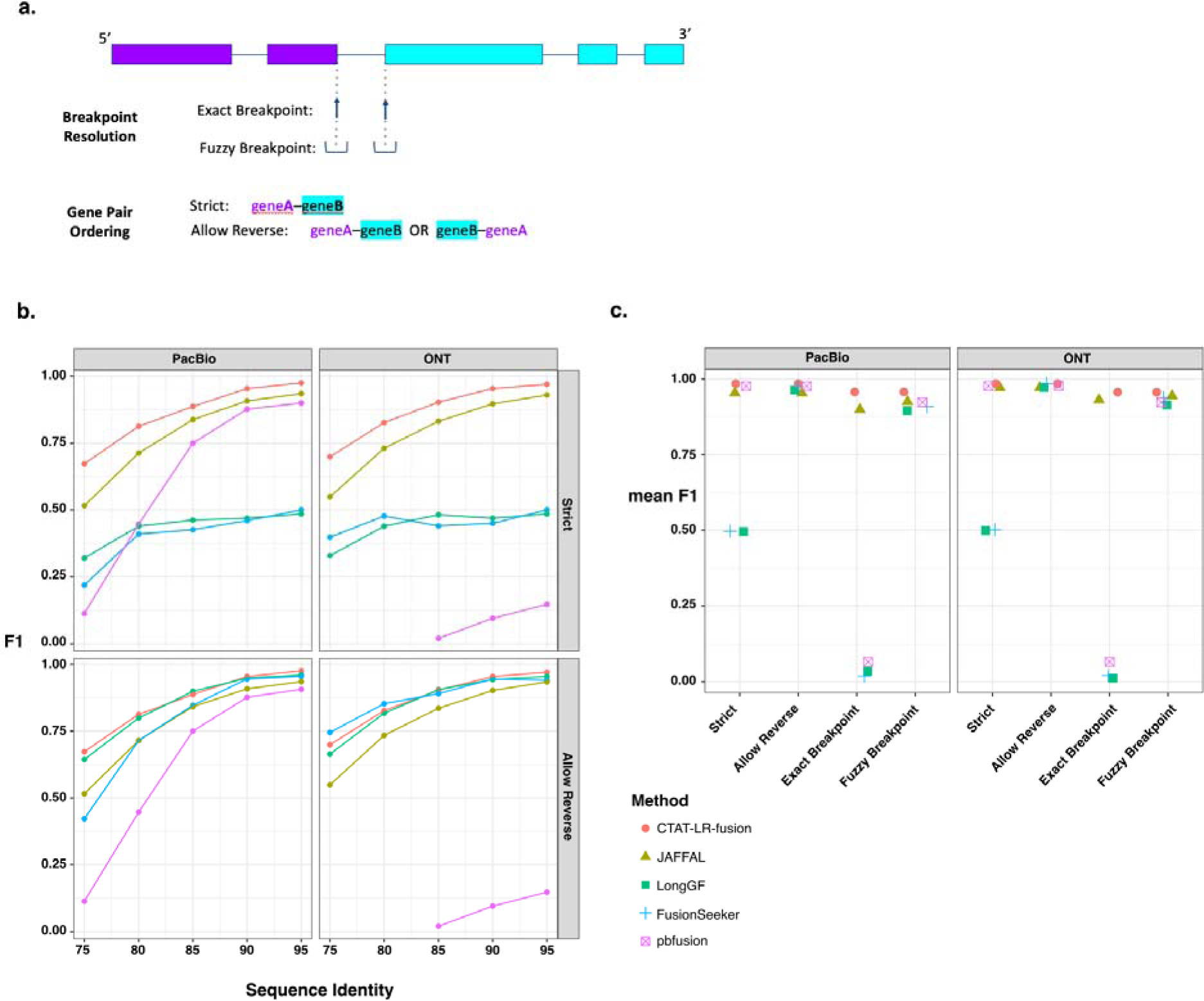
Accuracy for fusion transcript detection using simulated long reads. (a) Scheme for criteria in benchmarking fusion detection. (b) Accuracy reported as maximum F1 score determined using simulated PacBio and ONT long reads with moderate to high error rates (test data derived from [Jaffal paper ref]. (c) Accuracy using pbsim3 simulated PacBio HiFi or ONT R10.4.1 isoform reads at 50x coverage additionally focused on breakpoint resolution, with mean of maximum F1 values across 5 samples of 500 different target fusions each.

While the above test data were useful to differentiate accuracy characteristics across methods, the sequence error rates do not reflect those of the currently available long read sequencing technologies, which have rapidly improved to now routinely yield long read sequences at 1% (Q20) to 0.1% error (Q30) or better (Marx 2023). To that end, we used PBSIM3 (Ono et al. 2022) to simulate PacBio HiFi and ONT R10.4.1 long reads and further investigated fusion transcript detection accuracy across methods. With these newly simulated reads, all methods demonstrated high fusion transcript detection accuracy when considering only the unordered pairs of genes. To further explore differences in accuracy characteristics of these methods, we evaluated their fusion transcript breakpoint detection accuracy (**Figure 2a,c**). In particular, we compared the known simulated fusion breakpoints to the chromosomal location of the estimated fusion transcript breakpoint at each gene for each method. Interestingly, similar to the fusion gene ordering, only CTAT-LR-fusion and JAFFAL demonstrated highly accurate fusion transcript breakpoint detection (ignoring gene ordering during breakpoint evaluation). While FusionSeeker, LongGF, and pbfusion demonstrated little capacity for detecting exact breakpoints, the vast majority of breakpoints they reported were within a short distance (+/-5 bases) from the ground truth breakpoints (**Figure 2c**).

### Long Read Fusion Isoform Detection with a Reference Fusion Control RNA Sample

To evaluate CTAT-LR-fusion with real transcriptome sequencing data, we leveraged a commercial reference RNA sample from SeraCare (Seraseq Fusion RNA Mix v4) containing a set of 16 clinically-relevant fusion transcripts mixed at a fixed concentration into a background of total RNA derived from a commonly used human cell line (GM24385). This reference RNA sample was sequenced for long reads using our newly developed MAS-ISO-seq method (Al’Khafaji et al. 2023) commercialized by PacBio as Kinnex for augmented sequencing throughput. Sequencing was performed in triplicate, with replicate-1 using MAS-ISO-seq in a monomeric format (similar to standard PacBio Iso-Seq) and replicates-2 and −3 using the standard MAS-ISO-seq 8-mer concatamer format (as in Kinnex). The higher sequencing depth (**Supplementary Table 1**) of the standard MAS-ISO-seq data sets yielded more long fusion reads than the monomer-based (Iso-Seq-like) library construction, but after normalization for sequencing depth, rate of recovery of fusion reads was roughly equivalent, consistent with the sequencing libraries being derived from the single sample (**Supp. Figure 1a,b**). For comparison of fusion detection with PacBio long isoform reads vs. Illumina short read RNA-seq, we further sequenced this SeraCare fusion reference standard using Illumina TruSeq as triplicate libraries with paired-end 151 base length reads. Both MAS-ISO-seq and TruSeq generated approximately 5M to 10M reads (or paired-end sequences for TruSeq) per replicate (**Supplementary Table 1**).

Before comparing fusion detection between long and short reads with the Seraseq fusion sequencing data, we first downsampled the PacBio MAS-ISO-seq reads to match total sequenced bases from the Illumina sequenced sample replicates, respectively. All 16 control fusions were detected by CTAT-LR-fusion across three downsampled replicates with a range of 2 to 52 long PacBio isoform reads per sample (**Figure 3a**). Although matched Illumina TruSeq RNA-seq was performed for each of three replicates and overall gene expression was significantly positively correlated between long and short read sequencing (**Supp. Figure 1c**), relatively few control fusion supporting reads were detected and not all fusions were detected across three replicates based on the Illumina short reads; all fusions were detected in at least one TruSeq replicate across all samples but were missing in at least one replicate for 9/16 control fusions based on FusionInspector (**Figure 3a**).

**Figure 3:**
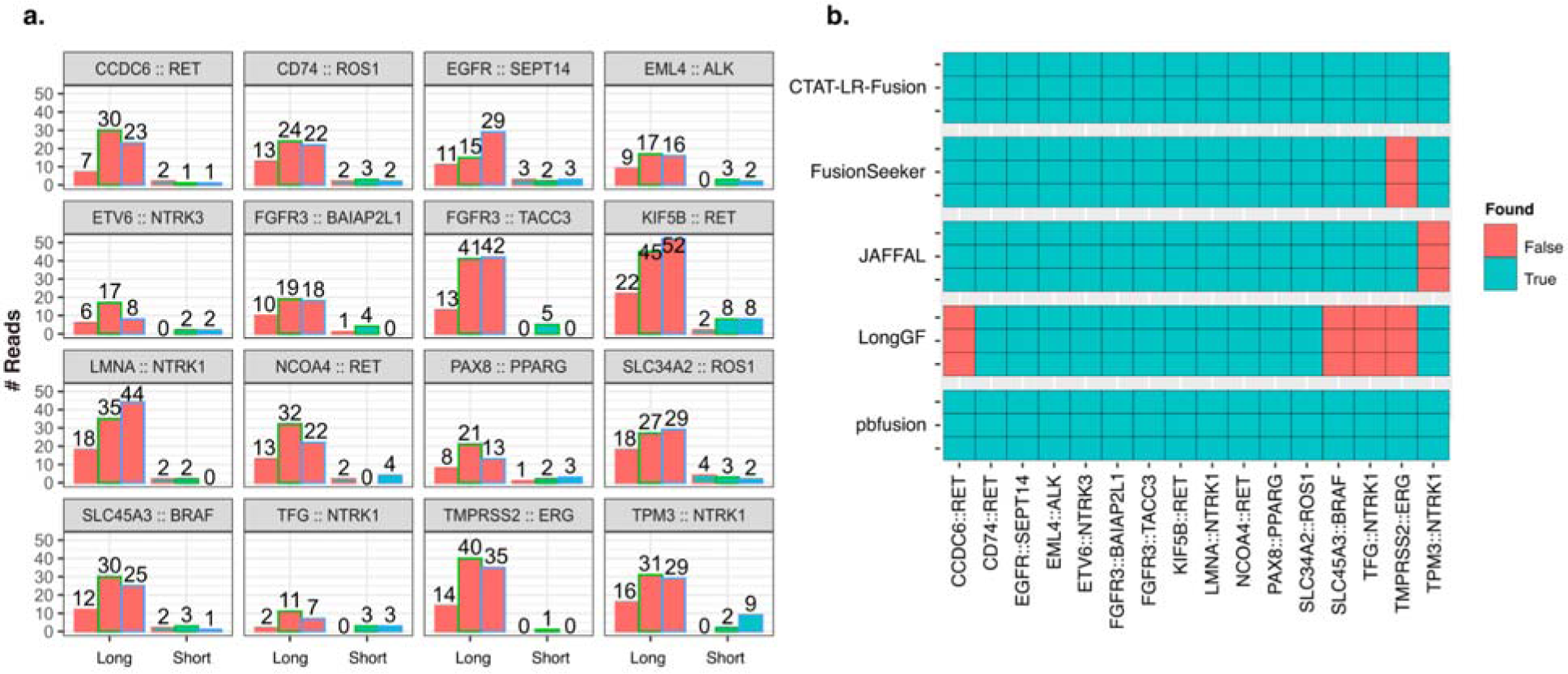
Fusion transcript detection applied to SeraCare v4 Fusion Reference Control sample. (a) Quantities of PacBio long reads and TruSeq Illumina short reads identified as evidence for each of the 16 control fusions as ascertained by CTAT-LR-fusion and FusionInspector, respectively, across each sample replicate. PacBio replicate reads were downsampled to match the number of sequenced bases from the respective Illumina replicate samples. (b) Binary heatmap for the identification of the 16 control fusions pairs in different fusion detection software according to each of the three replicates of long read sequences, using all (not downsampled) sequenced reads. PacBio replicates are ordered (a) left to right or (b) top to bottom as MAS-ISO-seq monomer (replicate 1), and MAS-ISO-seq 8mer-concatamer sequenced replicates 2 and 3. Counts of sequenced reads are provided in **Supplementary Table S1**.

We examined the alternative long read fusion transcript detection methods for identification of the 16 control fusions using all PacBio sequenced long isoform reads (**Figure 3b**). Only CTAT-LR-fusion and pbfusion (as of v0.4.0) were found to identify each of the 16 control fusions across each of the three long read sequencing libraries. Fusionseeker and JAFFAL each failed to report one of the 16 fusions, each a different fusion and consistent across all replicates.

LongGF, while having high accuracy for detection of fusions with simulated data, surprisingly was found least effective here in consistently missing 4/16 control fusions, only one of which was missed in common with another method: TMPRSS2::ERG, the hallmark fusion of prostate cancer, missed by both LongGF and FusionSeeker, while CTAT-LR-fusion detects 45, 98, and 104 long isoform reads supporting TMPRSS2::ERG across the three sequenced libraries.

### Long Read Fusion Isoform Detection from MAS-ISO-seq of Nine Cancer Cell Lines

We further explored long read based fusion transcript detection using transcriptomes from nine cancer cell lines derived from diverse cancer types including breast cancer (SKBR3, HCC1187, HCC1395), prostate cancer (VCaP), chronic myelogenous leukemia (K562), ALK+ anaplastic large cell lymphoma (KIJK), T cell lymphoma (MJ), small cell lung cancer (DMS53), and urothelial bladder cancer (RT112). Several of these cell lines are known to harbor oncogenic fusions including BCR::ABL1 in K562, TMPRSS2::ERG in VCaP, NPM1::ALK in KIJK, and FGFR3::TACC1 in RT112. We sequenced the transcriptomes of each cell line using PacBio MAS-ISO-seq (∼3-6M reads per sample, **Supplementary Table 1**) and called fusions using each long read fusion transcript prediction method (**Supplementary Table 2**). Counts of fusions predicted by each method having at least three long isoform reads as evidence vary greatly by cell line and by method, with RT112 and KIJK having the fewest fusion predictions, VCaP having the most, and the FusionSeeker method producing the greatest numbers of fusion predictions across all cell lines (**Figure 4a**). Altogether, we find 133 fusions agreed upon by at least two long read fusion prediction methods, with as few as 3 identified in cell line MJ and as many as 31 in VCaP (**Figure 4a**). Eight COSMIC fusions with known relevance to cancer biology including the hallmark fusions mentioned above were identified among most (6/9) of the cell lines and identified by at least two prediction methods with similar quantities of reads for each fusion, spanning two orders of magnitude (2 reads for K562|BCR::ABL1 to 463 reads for KIJK|ALK::NPM1)(**Figure 4b**).

**Figure 4.**
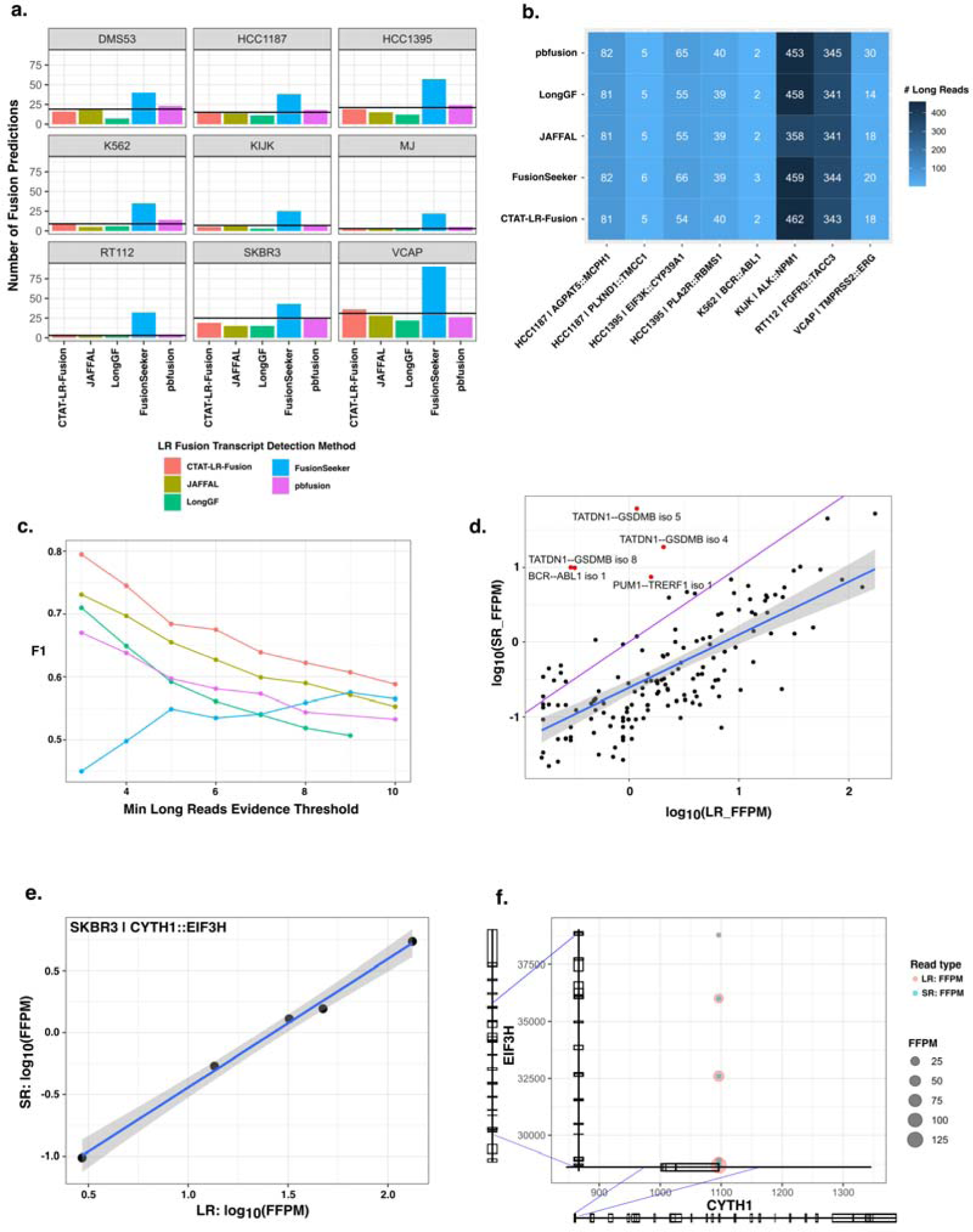
Detection of fusion transcripts from MAS-ISO-seq of 9 cancer cell lines. (a) Counts of fusion predictions according to cell line and prediction method, requiring a minimum of 3 long reads as supporting evidence. Line drawn indicates the number of fusions agreed upon by at least two methods. (b) Numbers of MAS-ISO-seq reads identified as evidence for COSMIC fusions according to method. (c) Fusion transcript detection accuracy according to minimum long reads supporting evidence based on the proxy truth set. (d) Comparison of long (MAS-ISO-seq) vs. short read (TruSeq Illumina) support for fusion isoforms detected by each according to CTAT-LR-fusion and FusionInspector, respectively. Read support is normalized for sequencing depth as FFPM. (e, f) Five fusion isoforms observed for the fusion gene CYTH1::EIF3H of cell line SKBR3 are (e) observed with highly correlated expression measurements as estimated from long and short RNA-seq reads and (f) shown according to fusion transcript breakpoints.

We separately sequenced these cell line transcriptomes using Illumina TruSeq with ∼30-50M paired-end 151 base length reads per sample (**Supplementary Table 1**), capturing read coverage across entire transcripts, and called fusions using STAR-Fusion. Of the 133 agreed-upon long read predicted fusions, more than half (79) were identified by STAR-Fusion with these short reads. Of another 354 fusions uniquely predicted from long reads by any method, only 12 (3%) were further identified using short reads.

Benchmarking fusion detection accuracy using these cell lines is challenging due to the lack of absolute truth sets, and experimental validations of fusions from these cell lines are not yet comprehensive. To assess accuracy, we employed a proxy truth set (as in (Haas et al. 2019)) where true fusions were operationally defined as those predicted by at least two different methods with at least 3 supporting reads, excluding likely artifacts and fusions with promiscuous fusion partners across samples, and treated uniquely predicted fusions as false positives (see **Methods**). We further incorporated the 12 Illumina-supported but otherwise uniquely predicted fusions along with the 133 agreed-upon fusion predictions as our proxy truth set. In benchmarking fusion detection for these cancer cell lines, CTAT-LR-fusion demonstrated superior performance across a range of minimum read evidence thresholds (**Figure 4c, Supp. Figure 2**). Only the performance of FusionSeeker was found to increase according to concomitant increase in required minimum read evidence support, primarily due to correspondingly large decreases of false positives (**Supp. Figure 2b**).

In exploring the fusion isoforms identified by CTAT-LR-fusion using combined long and short reads we found 213 fusion genes with 288 fusion splicing isoforms having both short and long read alignments together supporting each of the fusion transcript breakpoints. Fusion expression evidence is significantly but moderately correlated between short and long reads (R=0.70, p<2.2e-16), and the fraction of fusion-supporting long reads tends to exceed the short reads, with notable exceptions (**Figure 4d, Supplementary Figure 3a**). Oncogenic driver fusion BCR::ABL1 is one notable outlier with >100-fold enrichment of short reads detecting the fusion breakpoint than long reads per GB sequenced, apparently due to the long length of the fusion transcript with the fusion breakpoint up to 5 kb from the very 3’ end of the fusion transcript and from where PacBio long read isoform sequencing initiates. Short read enrichment for fusion detection was observed as weakly but significantly correlated (R=0.28, p=2.6e-8) with distance from the 3’ end of the fusion transcript (**Supplementary Figure 3b**).

Seven fusion genes were found with at least three fusion splicing isoforms each, including CYTH1::EIF3H in cell line SKBR3 with five alternatively spliced fusion isoforms with near perfectly positively correlated fusion expression as measured from long or short reads (R=0.997, p=1.9e-4, **Figure 4e,f**). The remaining examples mostly involved lowly expressed fusions with weakly-or un-correlated expression as measured according to short and long read support (**Supplementary Figure 4a**). Among these multi-isoform fusions, having access to both long and short reads yielded evidence for fusion isoforms uniquely supported by each read type. For example, TMPRSS2::ERG in VCaP has evidence for five fusion splicing isoforms where one is solely supported by long reads (**Supplementary Figure 4b**). In contrast, fusion TATDN1::GSDMB in SKBR3 has evidence for 13 fusion splicing isoforms, four of which are supported uniquely by short reads (**Supplementary Figure 4c**).

### Long Read Fusion Isoform Detection from Tumor Single Cell Transcriptomes

To examine CTAT-LR-fusion and long read isoform sequencing for fusion transcript detection in single cells, we leveraged earlier published PacBio single cell isoform sequencing data from two recently published studies: a T-cell infiltrated melanoma tumor sample from (Al’Khafaji et al. 2023), and three different metastatic high grade serous ovarian carcinoma (HGSOC) omental samples from (Dondi et al. 2023). In both studies, matching sample Illumina RNA-seq data was available, enabling us to further explore differences in detection of fusion transcripts based on long vs. short read sequencing. In these single cell applications, the 10x Genomics single cell sequencing libraries were based on 3’ end sequencing, inherently biasing sequencing coverage to the very 3’ ends of sequenced isoforms with Illumina RNA-seq.

The sequenced T-cell infiltrated melanoma tumor sample consisted of 6932 cells including 701 tumor cells (10%), sequenced with 21M PacBio MAS-ISO-seq reads and 207M single-end 55 base length reads (**Supplementary Table 1**). Fusion transcripts were examined using CTAT-LR-fusion for PacBio long reads and STAR-Fusion and FusionInspector for Illumina short reads (**Supplementary Table 3**). Only one fusion was found in more than 1% of tumor or normal cells: NUTM2A-AS1::RP11-203L2.4 found in 265 tumor cells (38%) and only 3 normal cells (0.05%) through a combination of long and short read fusion transcript analyses (**Figure 5a**); only short read fusion evidence was found corresponding to these 3 normal cells, all 3 detected by FusionInspector and one by STAR-Fusion, and such reads might have derived from ambient tumor RNA. Approximately 60% of the NUTM2A-AS1::RP11-203L2.4 containing tumor cells were solely identified by long read evidence, another 20% by short reads only, and the remaining 20% by both short and long reads (**Figure 5b**). Interestingly, fusion gene partner NUTM2A-AS1 has recently been identified as an oncogene with roles in multiple cancer types (Wang et al. 2020; Wang et al. 2021; Long et al. 2023). The long fusion reads appear to be largely full-length and yield evidence for eight different fusion splicing isoforms, mostly involving skipping of alternative exons and one isoform involving an alternative terminal exon (**Figure 5c**). The short read alignments provide evidence for five alternatively spliced isoforms but because of the short read length only the partial isoform structure around the fusion transcript breakpoints were resolved as opposed to the complete isoform structures clearly evident from the long reads (**Figure 5c**).

**Figure 5:**
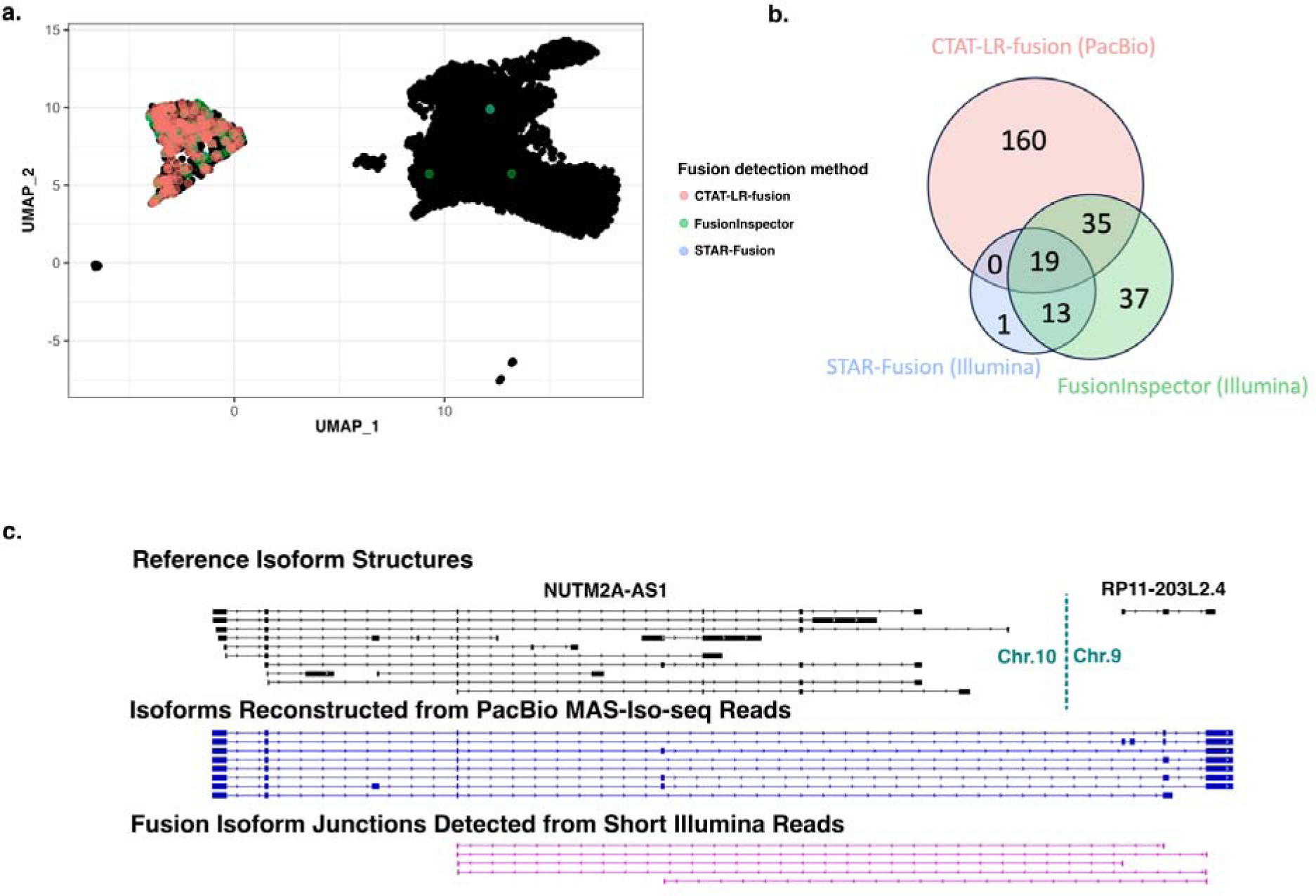
Detection of Fusion NUTM2A-AS1::RP11-203L2.4 in a T-cell infiltrated melanoma tumor sample. MAS-ISO-seq and matched Illumina RNA-seq data from a melanoma tumor sample M132TS 10x single cell library [published in (Al’Khafaji et al. 2023) were examined for fusion transcripts using CTAT-LR-fusion for PacBio long reads and STAR-Fusion and FusionInspector for Illumina short reads. (A) UMAP for melanoma sample M132TS single cells. Cells identified with the NUTM2A-AS1::RP11-203L2.4 fusion transcript are colored according to the detection method, predominantly labeling the cluster of malignant cells. (B) Venn diagram indicating the numbers of fusion-containing cells according to detection methods. (C) Fusion supporting read alignments and derived transcript isoform structures based on long (center) or short (bottom) read sequences in the context of the FusionInspector modeled gene fusion contig. Gencode v22 reference isoform transcript structures for NUTM2A-AS1 and RP11-203L2.4 genes are shown at top.

We explored the PacBio long isoform reads and Illumina short reads available for three HGSOC patient samples sequenced at single cell resolution. Here, tumor samples were derived from omental metastases, and for Patients 1 and 3, matched normal omentum samples were similarly processed and analyzed for comparison (all fusion predictions available as **Supplementary Table 4**). Numbers of PacBio long reads ranged from 22-54M reads along with matched 35-102M Illumina 56 base length single-end reads (**Supplementary Table 1**). In addition to identifying previously described fusions for these samples, we identified additional fusion genes and fusion isoforms supported by long and/or short RNA-seq reads, with multiple different fusion gene products generated from the same genome restructuring events. For detecting somatic cancer-specific fusions in these samples, we required at least five tumor cells to exhibit long or short read RNA-seq alignment evidence, and for identified fusions to be missing from matched normal samples where available.

Sequencing of the Patient-1 tumor sample yielded 497 total cells, with 92 cells (19%) identified as HGSOC cells, from which we identified only four somatic fusion transcripts: SMG7::CH507-513H4.1 (26 cells), RAPGEF5—AGMO (6 cells), NTN1--CDRT15P2 (5 cells), and GS1-279B7.2--GNG4 (5 cells) (**Supplementary Table S5**). For RAPGEF5::AGMO, half (3/6) of the cells were detected only by long reads, and 1/6 exclusively by short reads. The other three fusions were found only by long reads. Expression-based clustering of cells for the Patient 1 tumor sample resolved two HGSOC cell clusters, with fusion RAPGEF5::AGMO evident in tumor cells largely clustered separately from cells expressing SMG7::CH507-513H4.1 and GS1-279B7.2--GNG4, potentially reflecting tumor heterogeneity **Figure 6a,b**). Fusion NTN1::CDRT15P2 was found expressed in both tumor cell clusters and more likely clonal (**Figure 6b**).

**Figure 6:**
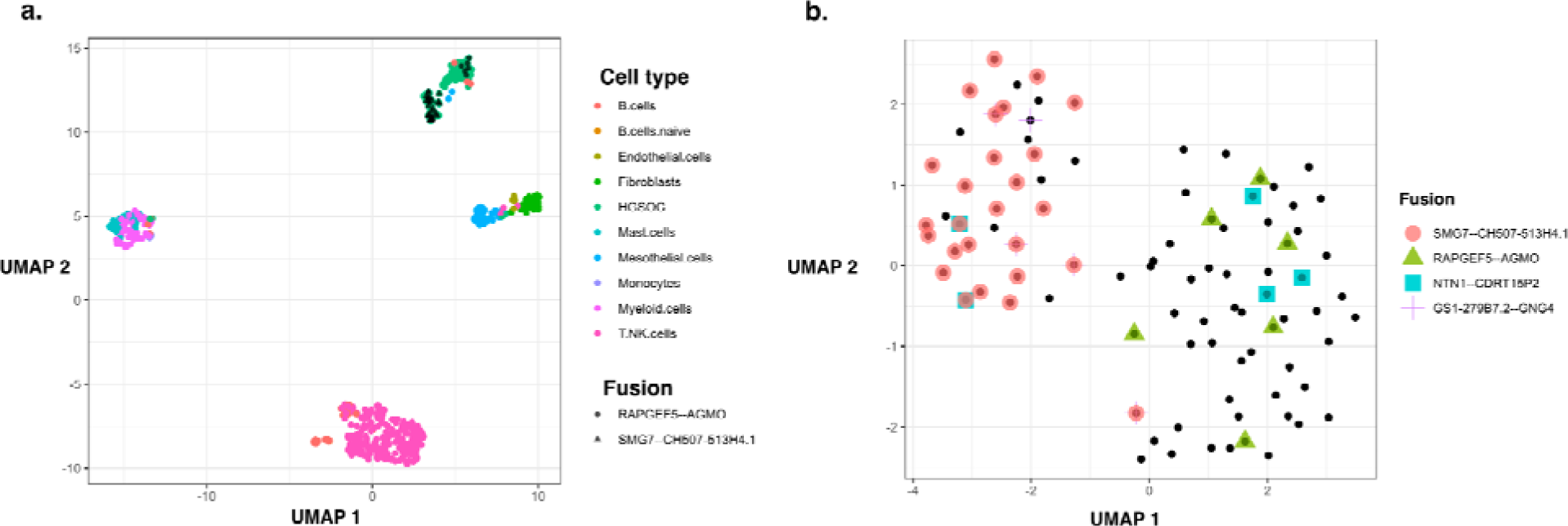
Fusion expression intra-tumor heterogeneity observed in cancer cells. (A) UMAP embedding of all cells from HGSOC Patient 1, colored by cell type. Fusion RAPGEF5::AGMO and SMG7::CH507-513H4.1 are expressed in two different HGSOC cell clusters. (B) UMAP embedding of HGSOC cells from HGSOC Patient 1, colored by fusions expressed. RAPGEF5::AGMO is expressed exclusively in the right cluster. SMG7::CH507-513H4.1 and GS1-279B7.2::GNG4 fusions coexpress and are expressed almost exclusively in the left cluster. The two NTN1::CDRT15P2 fusion expressing cells in the left cluster co-express the SMG7::CH507-513H4.1 fusion.

The Patient-2 tumor sample yielded 453 total cells, with 208 (46%) identified as HGSOC cells, from which we identified 16 different malignant cell enriched fusion transcripts (**Supplementary Table S5**), including the earlier-identified IGF2BP2::TESPA1 fusion between chr3 and chr12 evident in 176/208 (85%) of the tumor cells. Another fusion is found with proximal breakpoints yielding fusion transcript SPATS2::TRA2B (21 tumor cells, 10%), and likely resulting from the same tumor genome rearrangements involving chr3 and chr12. Both of these fusions were detected via long and short RNA-seq reads. While a single fusion splicing isoform dominated IGF2BP2::TESPA1 detection in cells by both long and short reads, additional fusion splicing isoforms were detected with only short read support according to both STAR-Fusion and FusionInspector (**Supplementary Table S4**). Nearly all (20/21) of the SPATS2::TRA2B expression cells are found to co-express IGF2BP2::TESPA1. Other notable fusions in the Patient 2 tumor sample involve known tumor oncogenes and include CBL::KMT2A (16 tumor cells) and DEK::CASC17(11 tumor cells), both identified solely by long reads. The previously reported FNTA fusion supported by long reads was missed here but manually verified, as the FNTA fusion partner transcribed region was lacking from the reference annotation and currently required for ctat-LR-fusion reporting. Another prevalent fusion PSMB7::SCAI (52 tumor cells) detected mostly by long reads and with four fusion splicing isoforms involves suppressor of cancer cell invasion gene SCAI. The reciprocal fusion SCAI::PSMB7 was previously detected in serous ovarian cancer cell line COV504_OVARY of the Cancer Cell Line Encyclopedia (Barretina et al. 2012), further implicating this rearrangement as of particular interest to this cancer type.

The Patient-3 tumor sample yielded 646 total cells with only 38 (6%) HGSOC cells. Here, only 2 fusions identified as enriched in the tumor cells: the previously identified CBLC::CTC-232P5.1 fusion in 16 cells and additionally found SNRNP70::ZIK1 in 8 cells (**Supplementary Tables S5**). Interestingly, each of these SNRNP70::ZIK1-expressing cells co-expressed the CBLC::CTC-232P5.1 fusion. Both fusions involve genes localized to the bottom arm of chr19 (CBLC and SNRNP70 transcriptional breakpoints within 5Mb), and potentially derive from the same genome restructuring events. There is evidence for five fusion transcript breakpoints for the CBLC::CTC-232P5.1 fusion indicating at least five fusion splicing isoforms, and all but one has support from both short and long reads. Fusion SNRNP70::ZIK1 was identified only by long reads.

Consistent with earlier studies, we find evidence of fusion transcripts expressed in normal cells, both from normal cells identified within the tumor microenvironment and from cells derived from the tumor-free matched normal samples. Excluding fusion transcripts previously identified in earlier large-scale studies of normal tissues, we find several fusion transcripts evident from the long isoform sequences that are patient-specific or in common across different patients, sometimes involving known oncogenes and previously implicated as potentially oncogenic.

Examples include fusion RP11-444D3.1::SOX5, previously implicated in endometrial cancer (Yao et al. 2019) and meningioma (Viaene et al. 2019) and recently reported as found in normal tissues in glioblastoma (Hernandez et al. 2022), but found here in small numbers of malignant (7) and normal (3) cells in the melanoma tumor sample and similarly identified among small numbers of cells (2 to 11) among each of the three HGSOC patient samples sets of tumor and matched normal samples. Fusion YWHAE::CRK involving fused oncogenes was detected in HGSOC Patient-1 normal sample in five mesothelial cells and in the tumor sample only one HGSOC cell. Fusion ZCCHC8--RSRC2, previously detected in several tumor studies (Yoshihara et al. 2015; Hu et al. 2018; Dehghannasiri et al. 2019; Jang et al. 2020; Haas et al. 2023), was identified as highly prevalent and broadly expressed across cell types in HGSOC Patient-3 tumor and matched normal samples, identified in 46% and 36% of sequenced cells, respectively.

## Discussion

As sequencing technologies and experimental methods continue to advance, we are faced with new challenges and opportunities for development of computational methods to extract deeper insights and further our understanding of biological systems. Rapid innovation in the long-read sequencing space has enabled full-length single cell RNA isoform sequencing, pushing the boundaries of transcriptome research. This leap in resolution has transformed our ability to accurately identify, discover, and quantify isoforms from genes and gene fusions, further accelerating biomedical research including studies of cancer and clinical applications to support personalized medicine.

Here we describe a new addition to our Trinity Cancer Transcriptome Analysis Toolkit (CTAT) for detection of fusion transcripts from long isoform read sequences called CTAT-LR-fusion. This module complements our earlier-developed Trinity CTAT methods available for detecting fusions based on shorter Illumina reads (usually 50-150 bases in length, single-end or paired-end), including TrinityFusion (Haas et al. 2019) for fusion transcripts based on genome-free Trinity (Grabherr et al. 2011; Haas et al. 2013) de novo assembled fusion isoforms, STAR-Fusion (Haas et al. 2019) for fusion detection based on chimeric short-read alignments, and FusionInspector (Haas et al. 2023) for supervised *in silico* validation of targeted gene fusions. Our CTAT-LR-fusion method for long isoform read fusion detection was motivated by TrinityFusion, using long isoform reads instead of Trinity-reconstructed transcripts for fusion detection, and by FusionInspector for modeling fusion gene contigs and quantification of fusion read support. FusionInspector is also further integrated into CTAT-LR-fusion as a submodule for evaluation of Illumina short read fusion evidence for candidates identified from the long reads in the case both long and short reads are provided as inputs.

We demonstrated superior accuracy of CTAT-LR-fusion for fusion detection based on long isoform reads derived from simulated data and from real data as derived from our application of high throughput PacBio long read RNA-seq, MAS-ISO-seq, to the Seraseq Fusion RNA Mix v4 control sample containing 16 spiked-in oncogenic fusion transcripts and to nine cancer cell lines. CTAT-LR-fusion was shown capable of robust identification of all 16 control fusions within the Seraseq fusion mix, and most accurate at identifying fusion transcripts based on simulated data across broad ranges of sequencing error. While high error rates are relegated to the earliest implementations of long read sequencing technologies, due to continued advancements in sequencing chemistries and computational methods for base-calling, contemporary sequencing accuracies of long reads no longer necessitate fusion detection methods compatible with high sequencing error rates. However, as newer and cheaper long read sequencing technologies are developed, the more extensive fusion detection capabilities of CTAT-LR-fusion could prove useful.

Proper detection and reporting of fusion transcripts require consideration of the order and orientation of the fused genes in the context of the fusion transcripts expressed and accurate reporting of the fusion transcript breakpoint, which most often involves standard transcript splicing that fuses an exon of one gene to an exon of the fusion partner. Of the evaluated long read isoform fusion detection methods, only CTAT-LR-fusion, JAFFAL, and pbfusion (as of v0.4.0) properly reported fusions in proper order and orientations along with precisely defined fusion isoform breakpoints. Reporting of fusion gene order and orientation is essential, as the alternate fusions made possible between two fusion genes have different interpretations and ramifications regarding oncogenicity, with relevance to clinical applications. For example, genes TACC3 and FGFR3 neighbor each other within a 100 kb region on chr4. A fusion detected as TACC3::FGFR3 could be considered an example of cis-splicing between neighboring genes, and potentially discarded as irrelevant. However, a genome rearrangement yielding the oncogenic fusion FGFR3::TACC3 (Costa et al. 2016) would be imperative to report. Other scenarios where fusion order and orientation are important considerations include reciprocal translocations, such as frequently encountered for the oncogenic BCR::ABL1 fusion among others (Haas et al. 2023). Finding BCR::ABL1 and its reciprocal ABL1::BCR fusions in the same patient sample via their distinct fusion transcripts could be considered evidence for a reciprocal chromosome translocation event. Note that in this case the BCR::ABL1 fusion transcript is the variant that yields the oncogenic fusion protein that drives tumorigenesis, and ABL1::BCR is likely collateral damage with questionable relevance to disease.

Accurate detection of fusion transcript breakpoints is essential for characterizing the splicing complexity of gene fusions. It is often the case that gene fusions produce multiple fusion transcript isoforms. For example, for fusion TATDN1::GSDMB in breast cancer cell line SKBR3, we find evidence of 13 distinct fusion transcript isoforms. Alternative splicing of fusion genes in cancer provides additional opportunities for neoantigen candidate discovery for applications in personalized immunotherapy, and their consideration could be especially useful when exploring cancers with low tumor mutation burden and limited candidates for neoantigen discovery based on expressed and translated somatic variants.

In all our applications of CTAT-LR-fusion to bulk and single cell transcriptomes presented here, we examined the capabilities of both long and short RNA-seq reads with matched samples. With few exceptions, fusion detection from long isoform reads greatly outperformed short reads, with more fusion genes and fusion transcript splicing isoforms and greater numbers of tumor single cells expressing fusions detected via long isoform reads. Perhaps unsurprisingly, fusion evidence is more concentrated among the long reads due to the sheer length of each long read, often providing full length isoform sequences for fused and normal isoforms of transcribed genes, as opposed to Illumina RNA-seq which entails fragmentation of long isoforms into shorter sequenceable fragments of transcripts, with fusion evidence restricted to the sequenced fragments of expressed transcripts. For single cell transcriptomes, the disparity between long and short reads widens as both long and short reads tend to initiate from the very 3’ end of transcripts. Detection of fusion isoforms based on short 3’ end sequences poses inherently strict limitations on short reads towards detecting breakpoints that occur proximal to the very 3’ end of the downstream fusion partner. In our survey of a melanoma tumor sample with single cell transcriptome data, long reads greatly outperformed short reads for detecting potentially oncogenic and tumor-specific NUTM2A-AS1::RP11-203L2.4 fusion-expressing cells. In our exploration of HGSOC tumor sample transcriptomes at single cell resolution, we mostly detected tumor-relevant fusions with long isoform reads.

Through combined use of short and long reads data, we increase detection sensitivity of gene fusions and numbers of cells with evidence of expressed fusions, demonstrating the synergy of both data types in bulk and single-cell samples. In bulk isoform sequencing, fractions of reads corresponding to fusion isoforms by long and short reads were significantly positively correlated, with specific examples such as CYTH1::EIF3H demonstrating near-perfect correlation.

Exceptions do exist where long or short reads were found to exclusively detect specific fusion isoforms or contrasting enrichments in detection of isoforms such that the dominant fusion splicing isoform detected via short reads was not always the dominant fusion isoform detected via long reads. Some differences such as the high enrichment of BCR::ABL1 fusion detection from short reads can be partially attributed to transcript breakpoints distal from the 3’ end and requiring very long isoform read sequencing to be able to traverse the breakpoint with long reads. Other differences are not yet understood and may reflect sequencing biases between platforms or sequencing protocols. As long read isoform sequencing becomes more routine, and as we explore increasing numbers of tumor cell lines and tumor single cell samples, we’ll have more opportunities to explore these differences, further optimize long read sequencing methods and continue to evaluate our toolkit and capabilities for integrated long and short RNA-seq along the way.

## Methods

### CTAT-LR-fusion long read fusion isoform detection

The CTAT-LR-fusion workflow has two phases: (1) initial rapid detection of fusion gene candidates and (2) fusion contig modeling with fusion candidate read alignment and breakpoint support quantification. These phases are described in detail below:

### CTAT-LR-fusion phase 1

Rapid detection of fusion gene candidates. Long isoform reads are aligned to the human reference genome using a customized version of minimap2 called ctat-minimap2 (https://github.com/TrinityCTAT/ctat-minimap2), which generates full read alignments only for reads that have preliminary mappings to multiple genomic regions. As most long reads are non-chimeric and mapped to single genomic regions, ctat-minimap2 avoids computational effort in generating alignments for reads that are unlikely to correspond to fusion genes, speeding up this initial read alignment stage 4-fold (see **Supplemental Code**). Chimeric read alignments derived from ctat-minimap2 are then assigned to reference gene annotations based on genomic coordinates. A preliminary list of fusion candidates is defined based on proximity to reference gene structures, requiring read alignments to have a default minimum of 70% alignment identity. Chimeric long reads are tallied according to candidate gene pairs and read alignment breakpoints are compared to the nearest neighboring exon boundaries. For all supporting reads, the minimum distance between exon boundaries and read alignment breakpoints are determined and candidate fusion gene pairs are pursued if either of the following conditions are met:

● Both chimeric alignment boundary minimum distances are within 50 bases of a reference transcript structure exon boundary.
● One chimeric boundary minimum distance is within 50 bases and the other is within 1kb of a reference transcript structure exon boundary, and multiple reads support the fusion between candidate gene pairs.

Fusion gene pair candidates are further filtered according to minimum expression threshold criteria (default: minimum 0.1 FFPM = at least 1 fusion long read per 10M total long reads), and such candidates are pursued in CTAT-LR-fusion phase 2 for further vetting and breakpoint quantification.

### CTAT-LR-fusion phase 2: Fusion contig modeling, long read realignment and breakpoint quantification

Phase 2 leverages techniques and methods in FusionInspector with modifications for long read alignment. Contig models for fusion genes are constructed using utilities in FusionInspector as previously described (Haas et al. 2023), positioning fusion gene structure candidates in the proposed order and orientation in single contigs with intronic regions shrunken to 1 kb. Candidate fusion-supporting long reads identified in Phase 1 are realigned to these fusion contigs using minimap2 (Li 2018). Read alignments with segments that terminate within 3 bases of a reference transcript exon boundary are snapped to that exon boundary, found useful for highly divergent read alignments and largely unnecessary for current HiFi reads. Fusion reads are identified as those that align spanning both genes in the fusion contig and breakpoints are tallied according to alignment ends that bridge the two genes. Fusions are filtered similarly as done for STAR-Fusion, requiring a minimum of 0.1 FFPM fusion expression evidence, and a minimum of 2 fusion reads where non-consensus splice dinucleotides exist at fusion breakpoints. By default, fusions known to occur in normal tissues are eliminated by looking up the GTEx fusions catalog, as incorporated into FusionAnnotator (Haas 2023) used with CTAT Human Fusion Lib (Haas 2021) (v0.3.0). Where there is evidence for multiple fusion splicing isoforms for a given fusion gene, those isoforms with less than 5% of the dominant isoform expression are discarded as potential noise.

When long reads are supplemented with Illumina short reads, FusionInspector is executed with the short reads and the fusion contig gene models derived from CTAT-LR-fusion Phase 1. The FusionInspector results are then merged with the CTAT-LR-fusion results based on long reads. In this case, filtering of fusion candidates is modified to consider results based on the short reads such that all fusion isoforms with a minimum of 0.1 FFPM as computed separately from long reads or short reads are included in the final report.

Fusion results based on single cell transcriptomes are further processed to generate per-cell fusion read support. Before running single cell transcriptome long or short reads through CTAT-LR-fusion, we encoded cell barcodes and read UMI data into the read name. The fusion reports from CTAT-LR-fusion and other CTAT fusion modules include lists of reads that support each fusion transcript isoform. From the read names in the fusion reports, we then extract the cell barcodes and UMIs and provide the per-cell reporting of fusion content.

### Fusion isoform detection via long read or short read sequencing

For each of the long read isoform sequencing based fusion prediction methods, we created docker images with the most recently available software versions installed. Workflows were built using WDL and data were processed using the Terra cloud computing framework. Software versions used are as follows: we used our latest CTAT-LR-fusion (v0.13.0) which we made available on GitHub at https://github.com/TrinityCTAT/CTAT-LR-fusion, JAFFAL (v2.3) from https://github.com/Oshlack/JAFFA, pbfusion (v0.4.0) from https://github.com/PacificBiosciences/pbfusion/releases, FusionSeeker (v1.0.1 commit 5710dc4 from https://github.com/Maggi-Chen/FusionSeeker, and LongGF(version 0.1.2) from https://github.com/WGLab/LongGF. Docker files and WDL workflows are made available at: https://github.com/broadinstitute/CTAT-LRF-Paper/tree/main/0.Workflows_and_Dockers. We prepared the reference data for each of the software based on its tutorial, and consistently used GRCh38 as the reference genome, and used GENCODE (Frankish et al. 2019) annotation version 22 for the transcriptome annotation. Illumina RNA-seq were analyzed using STAR-Fusion v2.12.0 and FusionInspector v2.8.0 as previously described (Haas et al. 2023).

### Simulated RNA-seq

Simulated fusion isoform reads were obtained from two sources: the JAFFAL published simulated data containing high error rates leveraging Badread (Wick 2019), and our own simulated high fidelity reads using PBSIM3 (Ono et al. 2022).

### Badread simulated fusion reads from the JAFFAL publication

We used the JAFFAL study (Davidson et al. 2022) simulated data for ONT and PacBio across the range of sequence divergences (75% identity to 95% identity), which was based on the set of simulated fusion transcripts sequences FASTA files generated in Haas et al, GB 2019 [31639029] for five different tissues (https://data.broadinstitute.org/Trinity/CTAT_FUSIONTRANS_BENCHMARKING/on_simulated_data/simulated_fusion_transcript_sequences/): adipose, brain, colon, heart, testis. The simulated JAFFAL datasets were downloaded from https://ndownloader.figshare.com/files/27676470.

### PBSIM3 simulated fusion reads

To reflect the error profiles of the latest PacBio and ONT sequencing technologies, we also simulated new ONT and PacBio long reads from these five different tissues using the long-read simulator PBSIM3 v3.0.1 (Ono et al. 2022) at 50x coverage as follows. To simulate PacBio HiFi reads, we first used PBSIM3 in full-length template-based mode (“--strategy templ”) with the provided PacBio Sequel continuous long reads (CLR) error model (“--errhmm data/ERRHMM-SEQUEL.model”) to generate multi-pass CLR sequencing data, producing 20 passes (“--pass-num 20”) for each input template to approximate high-accuracy HiFi reads; and then ran the PacBio CCS program v6.4.0 (https://github.com/PacificBiosciences/ccs) to generate HiFi reads from the multi-pass sequencing data produced by PBSIM3. To simulate ONT R10.4.1 reads, we similarly used the PBSIM3 full-length template-based simulation mode (“--strategy templ”) and the recently provided error model trained on R10.4 data (“--errhmm data/ERRHMM-ONT-HQ.model”) with a mean accuracy of 98% (“--accuracy-mean 0.98”), as recommended by PBSIM3 authors for ONT R10.4.1 reads (https://github.com/yukiteruono/pbsim3/issues/12). To obtain the desired coverage, we created multiple copies of the initial tissue templates and provided the resulting FASTA file as the “--template” parameter to PBSIM3. To link the reads to the original templates from which they were simulated for benchmarking, we made a small update to the PBSIM3 code in a PBSIM3 fork (https://github.com/MethodsDev/pbsim3) to report the read to template name mapping.

### Benchmarking of fusion transcript detection

When benchmarking using simulated long read fusion sequences, we parsed the gold standard fusion genes and breakpoints from sequences names in the simulated fusion transcripts sequence FASTA files (See **Simulated RNA-seq** section above).

We assessed the true positive (TPs), false positive (FPs) and false negative (FNs) for each fusion detection method by comparing their predictions against the respectively defined truth set. To quantify and compare the fusion detection performance, we applied three standard metrics for benchmarking fusion detection:

1. precision = TP / (TP+FP)
2. recall = TP / (TP+FN)
3. F1 = 2*precision*recall / (precision + recall)

For fusion genes, we have two modes of benchmarking by defining different levels of properly true positives: strict and “allow reverse”. In strict mode, we compared both of the gene pairs while strictly keeping their predicted gene order geneA::geneB, and assessed each fusion by matching both pairs of the genes with their official gene symbols, gene symbols for paralogs, and genes with overlapping coordinates along the genome. In “allow reverse” mode, we allowed the predicted gene order to be geneA::geneB or geneB::geneA when comparing with the corresponding truth set. For both geneA and geneB, gene symbols for genes with overlapping genomic coordinates were allowed as proxies and scored equivalently.

For breakpoints comparisons, we also implemented fuzzy or exact modes of performing the benchmarking. The two breakpoints were always sorted before comparison in either mode. In exact mode we strictly compared the sorted two breakpoint genomic coordinates for identity, and in fuzzy mode we expanded the allowed breakpoints of a fusion event to a window encompassing 5 bases upstream and downstream from each breakpoint.

When benchmarking using bulk cancer cell lines MAS-ISO-seq data, we filtered all the methods fusion calls based on 3 minimum long reads support. We further excluded fusions that tend to be enriched for artifacts, commonly encountered fusion from normal samples, or likely resulting from cis-splicing of neighboring transcripts; specifically, we filtered fusions including mitochondrial genes, HLA genes, gene pairs involving immunoglobulin gene rearrangements, fusions involving neighboring genes within 100 kb on a chromosome, or any fusions annotated as previously found in normal samples according to FusionAnnotator. Fusions passing these criteria were further filtered to retain fusions most relevant to individual cell lines by excluding fusions that involved promiscuous genes reported in fusion predictions by at least two different methods across at least three of the nine different cell lines examined here. After filtering, we defined truth set (TPs) as those fusions predicted by at least two different predictors, and FPs as fusions uniquely predicted by the corresponding method. Precision, recall, and F1 metrics were computed using this truth set. We examined how accuracy changed as a function of strength of evidence by evaluating accuracy metrics after filtering fusion predictions according to minimum read support (eg. **Supp. Figure 2a**).

A small fraction of pbfusion v0.4.0 results (∼1%) involved complex fusions involving multiple partners that were not always clearly identified with breakpoint information. For benchmarking purposes, we ignored instances where there lacked a clear one-to-one mapping between breakpoint coordinates and fusion partners, as recommended by the pbfusion developers (personal communication). In evaluation of the SeraCare fusions, the pbfusion output was manually examined to confirm capture of a reference fusion where breakpoint information was not clearly defined.

All benchmarking analysis code and the raw outputs from each of the evaluated prediction methods are available at: https://github.com/fusiontranscripts/LR-FusionBenchmarking.

### Bulk 8-mer MAS-ISO-seq for nine DepMap cell lines and two SeraCare fusion mix v4 replicates

#### RNA QC of Cancer Cell lines and Seraseq Fusion RNA mix

RNA samples were extracted form 9 cancer cell lines (VCAP, MJ, K562, RT112, KIJK, HCC1187, HCC1395, DMS53, and SKBR3) using Qiagen’s RNEasy Plus Kit (Qiagen, cat. no. 74134), and RNA from the Seraseq Fusion RNA mix v4 (SeraCare, cat. no. 0710-0497) were quality checked using a High Sensitivity RNA ScreenTape (Agilent, cat. no’s. 5067-5579 and 5067-5580) on an Agilent 4150 TapeStation system (Agilent, cat. no. G2992AA) to determine RNA Integrity Number (RIN) prior to first strand synthesis (FSS).

#### cDNA Synthesis from Cancer Cell Lines and SeraCare Fusion RNA mix

For both the cancer cell lines and the Seraseq Fusion RNA mix, cDNA was generated from RNA using components from a NEBNext® Single Cell/Low Input cDNA Synthesis & Amplification Module (New England Biolabs, cat. no. E6421S). The RNA Samples were diluted, the cancer cell lines to 50 ng/µl, and the SeraSeq fusion RNA mix to 15ng/ul. Per sample, the diluted RNA (200ng/cancer cell line sample, 100ng/SeraSeq fusion mix) was combined with 3µL of water, and 2µL of NEBNext Single cell RT primer (Sequence: AAG CAG TGG TAT CAA CGC AGA GTA CTT TTT TTT TTT TTT TTT TTT TTT TTT TTT TV), mixed via pipetting, and incubated at 70° C for 45 minutes before cooling to 20° C. Each reaction was then immediately combined with a second reaction mix consisting of 5µl of NEBNext Single Cell buffer, 2µl of NEBNext Single Cell RT Enzyme Mix, and 3µl of Nuclease-free water. The reaction was then incubated at 42°C for 45 minutes before being removed from the thermal cycler, having 1µl of 100µM Template switch oligo (Sequence; GCA ATG AAG TCG CAG GGT TrGrG rG) mixed in via pipetting, returning the reaction mix to the thermal cycler and incubating at 42°C for 15 minutes, then 85°C for 5 minutes, holding at 4°C. 30µl of elution buffer was added to each reaction for a total volume of 50µl, each reaction was then cleaned using 40µL (0.8x reaction volume) of SPRI beads (Beckman Coulter Inc, B23318) according to the manufacturer’s recommendations. The reaction was eluted in 50µl of elution buffer. 15µl of each cDNA was taken from the previous elution volume, and then combined with 25µl of NEBNext Single Cell cDNA PCR Master Mix,

2.5µl of 5µM Forward Primer (Sequence: AAG CAG TGG TAT CAA CGC AGA G), 2.5µl of an Indexed reverse primer (Sequence, variable, see **Supplementary Table S6**) and 5µl of Nuclease-free water for a total volume of 50µl. The reaction was mixed and then incubated in the thermal cycler for one cycle of 3 minutes at 98°C, 12 cycles of 20 seconds at 98°C – 30 seconds at 62°C – 8 minutes at 72°C, then one cycle of 5 minutes at 72°C, holding at 4°C. Each reaction was then cleaned using 35µL (0.7x reaction volume) of SPRI beads. The reaction was eluted off the beads in 50µl of elution buffer. The samples were quantified using a Qubit Flex Fluorometer (Thermo Fisher Scientific, cat. no. Q33327) and Qubit dsDNA HS Assay kit (Thermo Fisher Scientific, cat. no. Q32854) and analyzed via High Sensitivity D5000 ScreenTape (Agilent, cat. no’s. 5067-5594, 5067-5593, and 5067-5592) on an Agilent 4150 TapeStation system. The resultant cDNA was diluted down to 5ng/µl.

#### PacBio SMRTBell library preparation

The following section of the sequencing preparation was completed using kit components from the MAS-Seq for 10x Single Cell 3’ kit (PacBio, cat. no. 102-659-600), as well as individually created oligos. A PCR master mix for each sample was made using 100µl of MAS PCR Mix, 20ng of cDNA in 4µl of volume, and 96µl of nuclease-free water for a total volume of 200µl. The master mix was mixed and 22.5µl aliquots were distributed to each well of a 0.2ml PCR tube strip (USA Scientific Inc., cat. no. 1402-2500) where a 2.5µl addition of a 5µM primer mix was added **(see Supplementary Table S7)**. The samples were mixed and incubated in the thermal cycler for an initial denaturation step of one cycle for 3 minutes at 98°C, then seven cycles of denaturation for 20 seconds at 98°C, annealing for 30 seconds at 68°C, and extension for 8 minutes at 72°C, finally, a terminal extension of one cycle for 5 minutes at 72°C, holding at 4°C.

After incubation, the entire volume of each strip tube was pooled into a 1.5ml tube (total volume 200µl) prior to a 0.95x SPRI bead clean. The resultant product was eluted into 50µl of elution buffer. The product was quantified via Qubit Flex Fluorometer. 47µl from the previous elution was transferred into a 0.2ml PCR tube, 10µl of MAS Enzyme was added to each reaction then pipette mixed. The reactions were then incubated for 30 minutes at 37°C, holding at 4°C. The reactions were removed, and two reaction mixes were added, the first consisted of 1.5µl of MAS Adapter A Fwd 1.5 µl of MAS Adapter Q Rev, and 20µl of MAS Ligation additive. The second reaction mix added consisted of 10µl of Mas Ligase Buffer, and 10ml of MAS Ligase for a total combined reaction of 100µl. The reaction was mixed with wide bore pipette tips (Mettler-Toledo Rainin LLC, cat. no. 30389241), prior to being incubated for 60 minutes at 42°C, holding at 4°C. The reactions were removed from the thermal cycler and 75µl (0.75x) of resuspended SPRI beads were added. The reactions were mixed thoroughly using wide bore pipette tips and then left to incubate at room temperature for 10 minutes. The reactions were placed on a magnetic strip to pellet the beads, which were then washed twice in 200µl of 80% ethanol. 45µL of elution buffer was added to the reactions after the second ethanol wash and were left to elute off the beads for five minutes at room temperature. The reaction was then added back on to the magnet and the 45µl eluted MAS Array was moved to a separate 0.2ml PCR tube. 42µL of each of the eluted MAS array was transferred to a new 0.2ml PCR tube and a reaction mix consisting of 6µl of Repair buffer, and 2µl of DNA Repair Mix, was added for a total volume of 50µl. The reaction was mixed using wide bore pipette tips before incubating for 30 minutes at 37°C, holding at 4°C. The reactions were removed from the thermal cycler and 37.5µl (0.75x) of resuspended SPRI beads were added, and then cleaned according to the manufacturer’s specifications. The reaction was eluted in 40µl of elution buffer. To the 40µl of eluted DNA, a reaction mix consisting of 5µl of Nuclease buffer and 5ml of Nuclease mix was added for a total volume of 50µl. The reaction was pipette mixed using wide bore pipettes then incubated for 60 minutes at 37°C, holding at 4°C. The reactions were removed from the thermal cycler and 37.5µl (0.75x) of resuspended SPRI beads were added. The reactions were mixed thoroughly using wide bore pipette tips and then left to incubate at room temperature for 10 minutes. The reactions were placed on a magnetic strip to pellet the beads, which were then washed twice in 200µl of 80% ethanol. 25µL of elution buffer was added to the reactions after the second ethanol wash and were left to elute off the beads for five minutes at room temperature. The reaction was then added back on to the magnet and the 25µl eluted MAS Array was moved to a separate 0.2ml PCR tube. The reaction was then quantified using a Qubit Flex Fluorometer, and characterized using a Genomic DNA ScreenTape Analysis (Agilent, cat. no’s. 5067-5366 and 5067-5365) on an Agilent 4150 TapeStation system.

### PacBio Monomeric MAS-ISO-seq for SeraCare fusion RNA mix v4

#### RNA QC of Seraseq Fusion RNA Mix v4 for Monomeric MAS-Seq

The RNA sample (Seraseq® Fusion RNA Mix v4, cat. no. 0710-0497) was quality checked using a High Sensitivity RNA ScreenTape(Agilent, cat. no’s. 5067-5579 and 5067-5580) on an Agilent 4150 TapeStation system (Agilent, cat. no. G2992AA) to determine RNA Integrity Number (RIN) prior to first strand synthesis (FSS).

#### cDNA Synthesis from Seraseq RNA Mix v4 for Monomeric MAS-Seq

cDNA was generated from RNA using components from a NEBNext® Single Cell/Low Input cDNA Synthesis & Amplification Module (New England Biolabs, cat. no. E6421S), MAS-Seq for 10x Single Cell 3’ kit (PacBio, cat. no. 102-659-600), and individually created oligos. The RNA mix was diluted to 10ng/µl and split iIto two separate reaction vessels. Per reaction, the diluted RNA (10ng/µl, 7µl total volume, 70 ng total) was combined with 2µL of NEBNext Single cell RT primer (Sequence: AAG CAG TGG TAT CAA CGC AGA GTA CTT TTT TTT TTT TTT TTT TTTTTT TTT TTT TV), mixed via pipetting, and incubated at 70° C for 45 minutes before cooling to 20° C. Each reaction was then immediately combined with a second reaction mix consisting of 5µl of NEBNext Single Cell buffer, 2µl of NEBNext Single Cell RT Enzyme Mix, and 3µl of Nuclease-free water. The reaction was then incubated at 42°C for 45 minutes before being removed from the thermal cycler, having 1µl of 100µM Template switch oligo (Sequence; GCA ATG AAG TCG CAG GGT TrGrG rG) mixed in via pipetting, returning the reaction mix to the thermal cycler and incubating at 42°C for 15 minutes, then 85°C for 5 minutes, holding at 4°C. 30µl of elution buffer was added to each reaction vessel for a total volume of 50µl, each reaction was then cleaned using 40µL (0.8x reaction volume) of SPRI beads (Beckman Coulter Inc, B23318) according to the manufacturer’s recommendations. The reaction was eluted off the beads in 50µl of elution buffer. 15µl of each cDNA reaction was aliquoted from the previous elution volume, and then combined with 25µl of NEBNext Single Cell cDNA PCR Master Mix, 2.5µl of MAS Capture Primer FWD (Sequence: AAG CAG TGG TAT CAA CGC AGA G), 2.5µl of MAS Capture Primer REV, and 5µl of Nuclease-free water for a total volume of 50µl. The reaction was mixed and then incubated in the thermal cycler for one cycle of 3 minutes at 98°C, 14 cycles of 20 seconds at 98°C – 30 seconds at 62°C – 8 minutes at 72°C, then one cycle of 5 minutes at 72°C, holding at 4°C. Each reaction was then cleaned using 35µL (0.7x reaction volume) of SPRI beads. The reaction was eluted off the beads in 50µl of elution buffer. The samples were quantified using a Qubit Flex Fluorometer (Thermo Fisher Scientific, cat. no. Q33327) and Qubit dsDNA HS Assay kit (Thermo Fisher Scientific, cat. no. Q32854) and analyzed via High Sensitivity D5000 ScreenTape (Agilent, cat. no’s. 5067-5594, 5067-5593, and 5067-5592) on an Agilent 4150 TapeStation system.

#### PacBio SMRTBell library preparation

The following section of the sequencing preparation was completed using kit components from the MAS-Seq for 10x Single Cell 3’ kit (PacBio, cat. no. 102-659-600), as well as individually created oligos. A PCR mix for the sample was made using 25µl of MAS PCR Mix, 5ng of cDNA in 2µl of volume, and 23µl of nuclease-free water for a total volume of 50µl. The master mix was mixed and a 45µl aliquot was distributed to one well of a 0.2ml PCR tube strip (USA Scientific Inc., cat. no. 1402-2500) where 5µl addition of a 5µM primer mix of primers A-FWD and Q-REV was added (A-FWD, Sequence: AGCTTACTUGTGAAGAUCTACACGACGCTCTTCCGATCT, Q-REV, Sequence:

AUGCACACAGCUACUAAGCAGTGGTATCAACGCAGAG). The sample was mixed and incubated in the thermal cycler for an initial denaturation step of one cycle for 3 minutes at 98°C, then seven cycles of denaturation for 20 seconds at 98°C, annealing for 30 seconds at 68°C, and extension for 8 minutes at 72°C, finally, a terminal extension of one cycle for 5 minutes at 72°C, holding at 4°C. After incubation, 47.5µl (0.95x) SPRI beads were added for a clean. The resultant product was eluted into 60µl of elution buffer. The product was quantified via Qubit Flex Fluorometer. 55µl was transferred into a 0.2ml PCR tube, 2µl of MAS Enzyme was added to each reaction then pipette mixed. The reaction was incubated for 30 minutes at 37°C, holding at 4°C. The reaction was removed, and two reaction mixes were added, the first consisted of 1.5µl of MAS Adapter A Fwd 1.5 µl of MAS Adapter Q Rev, and 20µl of MAS Ligation additive. The second reaction mix added consisted of 10µl of Mas Ligase Buffer, and 10ml of MAS Ligase for a total combined reaction of 100µl. The reaction was mixed with wide bore pipette tips (Mettler-Toledo Rainin LLC, cat. no. 30389241), prior to being incubated for 60 minutes at 42°C, holding at 4°C. The reactions were removed from the thermal cycler and 75µl (0.75x) of resuspended SPRI beads were added and cleaned according to the manufacturer’s recommendations. The reaction was eluted in 45µl of elution buffer 42µL of the eluted MAS array was transferred to a new 0.2ml PCR tube and a reaction mix consisting of 6µl of Repair buffer, and 2µl of DNA Repair Mix was added for a total volume of 50µl. The reaction was mixed using wide bore pipette tips before incubating for 30 minutes at 37°C, holding at 4°C. The reactions were removed from the thermal cycler and 37.5µl (0.75x) of resuspended SPRI beads were added, and then cleaned according to the manufacturer’s recommendations. The reaction was eluted in 40µl of elution buffer. To the 40µl of eluted DNA, a reaction mix consisting of 5µl of Nuclease buffer and 5ml of Nuclease mix was added for a total volume of 50µl. The reaction was pipette mixed using wide bore pipettes then incubated for 60 minutes at 37°C, holding at 4°C. The reactions were removed from the thermal cycler and 37.5µl (0.75x) of resuspended SPRI beads were added and cleaned according to the manufacturer’s recommendations. The reaction was eluted in 25µl of elution buffer. The final product was then quantified using a Qubit Flex Fluorometer and characterized using a High Sensitivity D5000 ScreenTape on an Agilent 4150 TapeStation system.

#### Illumina TruSeq RNA-seq for nine DepMap cell lines and three SeraCare fusion RNA mix v4 replicates

DepMap samples were quantified by Qubit Ribogreen and normalized to 350 ng inputs respectively for the TruSeq stranded RNA protocol. All samples were determined by Agilent BioAnalyzer to have high quality with RINS > 9. Poly-adenylated RNAs were selected prior to fragmentation on the Covaris. Stranded cDNA libraries were generated following the Illumina TruSeq Stranded Total RNA protocol (<u>TruSeq Stranded Total RNA Reference Guide</u>). cDNA libraries incorporating ligated adapters were pooled and loaded on the NovaSeq SP for paired-end 151 bp sequencing targeting 50M paired reads per sample.

#### Single cell RNA-seq data

Melanoma sample M132TS – used previously published data from Aziz et al. (Al’Khafaji et al. 2023). This earlier publication focused on the T-cells and here we focused on the tumor cells, and so we extracted both and reprocessed through CellBender (Fleming et al. 2023). HGSOC – used previously published data from Dondi et al. (Dondi et al. 2023), reads downloaded from the European Genome-Phenome Archive (EGA) (Freeberg et al. 2022) under accessions EGAD00001009814 (PacBio) and EGAD00001009815 (Illumina). Cell annotations and long read gene counts per cell were retrieved from Dondi et al. For visualization, counts were normalized independently for each patient using sctransform (Hafemeister and Satija 2019), regressing out cell cycle effects and library size as non-regularized dependent variables. Similar cells were grouped using Seurat FindClusters (Satija et al. 2015). The results of cell clustering and cell typing were visualized in a low-dimensional representation using Uniform Manifold Approximation and Projection (UMAP) (Leland McInnes 2018).

## Supplemental Code

All analyses and figures generated as part of this work are available at https://github.com/broadinstitute/CTAT-LRF-Paper.

## Data Access

Simulated fusion reads leveraged from the earlier JAFFAL study (Davidson et al. 2022) were downloaded from https://ndownloader.figshare.com/files/27676470. Our PBSIM3 simulated fusion reads are available at Zenodo at: https://zenodo.org/records/10650516 doi:10.5281/zenodo.10650516. Illumina TruSeq and PacBio MAS-ISO-seq reads generated for the SeraCare SeraSeq Fusion Mix RNA v4 are available in SRA under BioProject ID PRJNA1076207, and for the nine DepMap cell line transcriptomes under BioProject ID PRJNA1077632. The human T-cell infiltrating melanoma single-cell RNA-sequencing data examined here and previously published in (Al’Khafaji et al. 2023) are available from dbGAP with accession number phs003200.v1.p1.

The HGSOC single cell data were obtained from EGA study EGAS00001006807 as data set IDs EGAD00001009814 (PacBio) and EGAD00001009815 (Illumina).

## Acknowledgements

We thank Simone Zhang, Andrew Tuttle, and Dev Gulati from the DepMap team who provided insight and assisted with the research; James Robinson and Helga Thorvaldsdottir for contributing and supporting igv-reports as used for our interactive reports of fusion-supporting read alignment evidence; Zev Kronenberg, Daniel Baker, and Roger Volden for addressing issues related to pbfusion; additional thanks to Z.K. for comments and suggestions regarding our manuscript; and Francis Jacob for his help through the HGSOC data access process. This work has been supported by National Cancer Institute grant U24CA180922 (B.J.H.), and partially funded by the Dependency Map Consortium. This work was supported by a Collaboration Agreement by and between Pacific Biosciences of California, Inc. and The Broad Institute, Inc. V.P. was supported by the Broad Institute Schmidt Fellowship. A.D. was supported by the European Union’s Horizon 2020 research and innovation program under the Marie Sklodowska-Curie grant agreement (#766030 to N. Beerenwinkel).

## Author Contributions

B.J.H. and Q.Q. wrote the initial manuscript draft, performed analyses, and contributed to CTAT-LR-fusion software development. B.J.H. and A.D. contributed to fusion discovery and analysis of the HGSOC single cell transcriptome data. K.W. prepared DepMap cell line RNA samples for sequencing. E.W. and A.S. contributed to sequencing of the SeraCare Seraseq Fusion Mix v4 RNA and the DepMap cell line RNA samples. A.K. contributed to processing of the short and long read RNA-seq to generate Fastq files used for downstream sequence analyses. V.P. contributed to the alignment optimization for chimeric reads and generated the PacBio and ONT synthetic benchmarking data. H.Y. contributed to processing and analysis of melanoma single cell transcriptome data. A.M.A. oversaw sample processing, sequencing, primary data processing and QC All authors contributed to the development of the final manuscript.

