## Supplementary Figures for "CTAT-LR-fusion: accurate fusion transcript identification from long and short read isoform sequencing at bulk or single cell resolution"

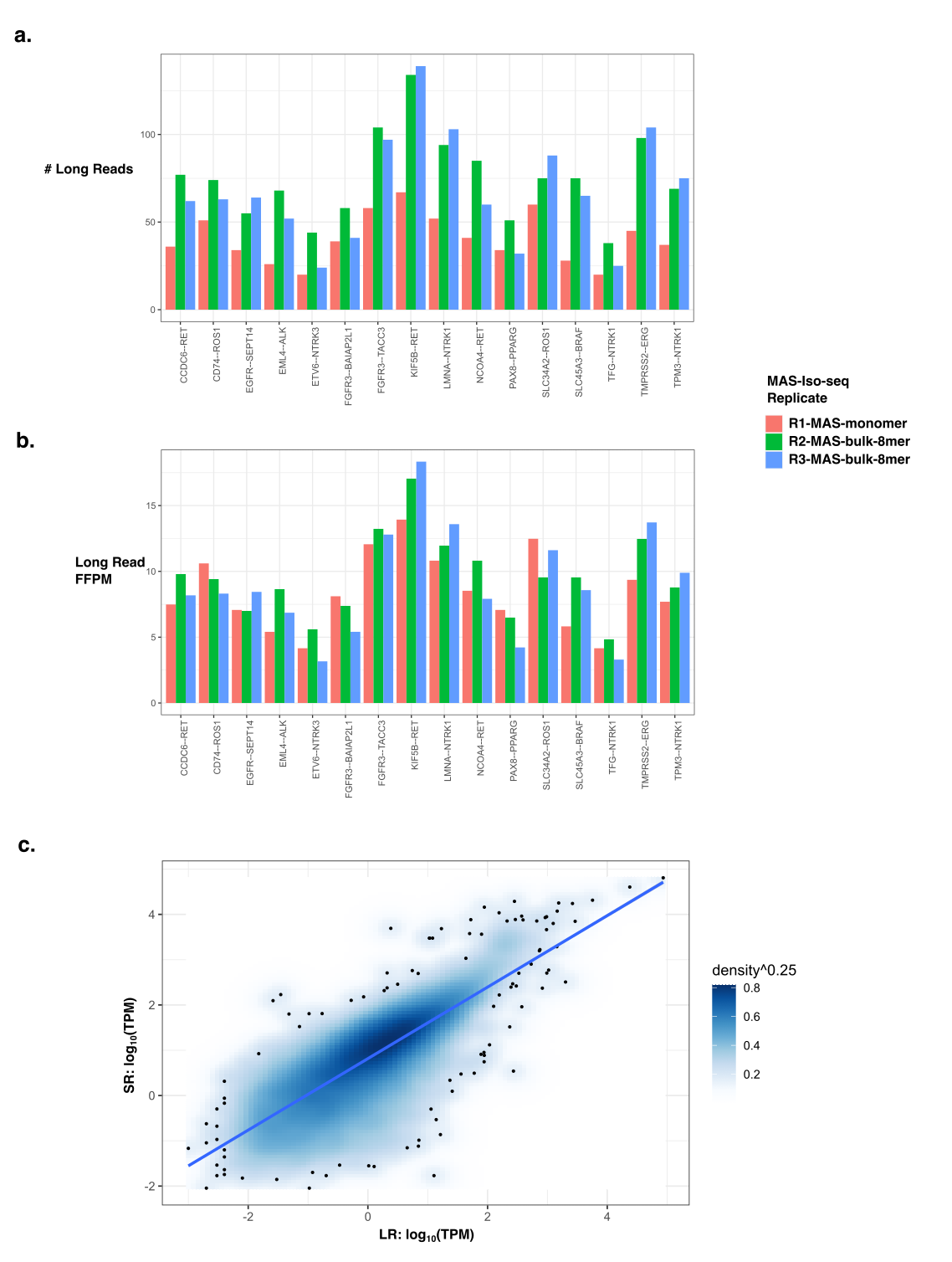

**Supplementary Figure 1: CTAT-LR-fusion detection of 16 control fusion transcripts based on PacBio long read isoform sequencing.** (a,top) raw counts, (b,bottom) normalized for sequencing depth as FFPM. (c) Gene expression quantification comparison based on long reads in comparison to short reads based on quantification of Gencode v22 reference annotations performed using Salmon. Expression values (TPM measurements) for PacBio long read sequencing and Illumina TruSeq short read sequencing are significantly positively correlated (R=0.79, p<2.2e-16).

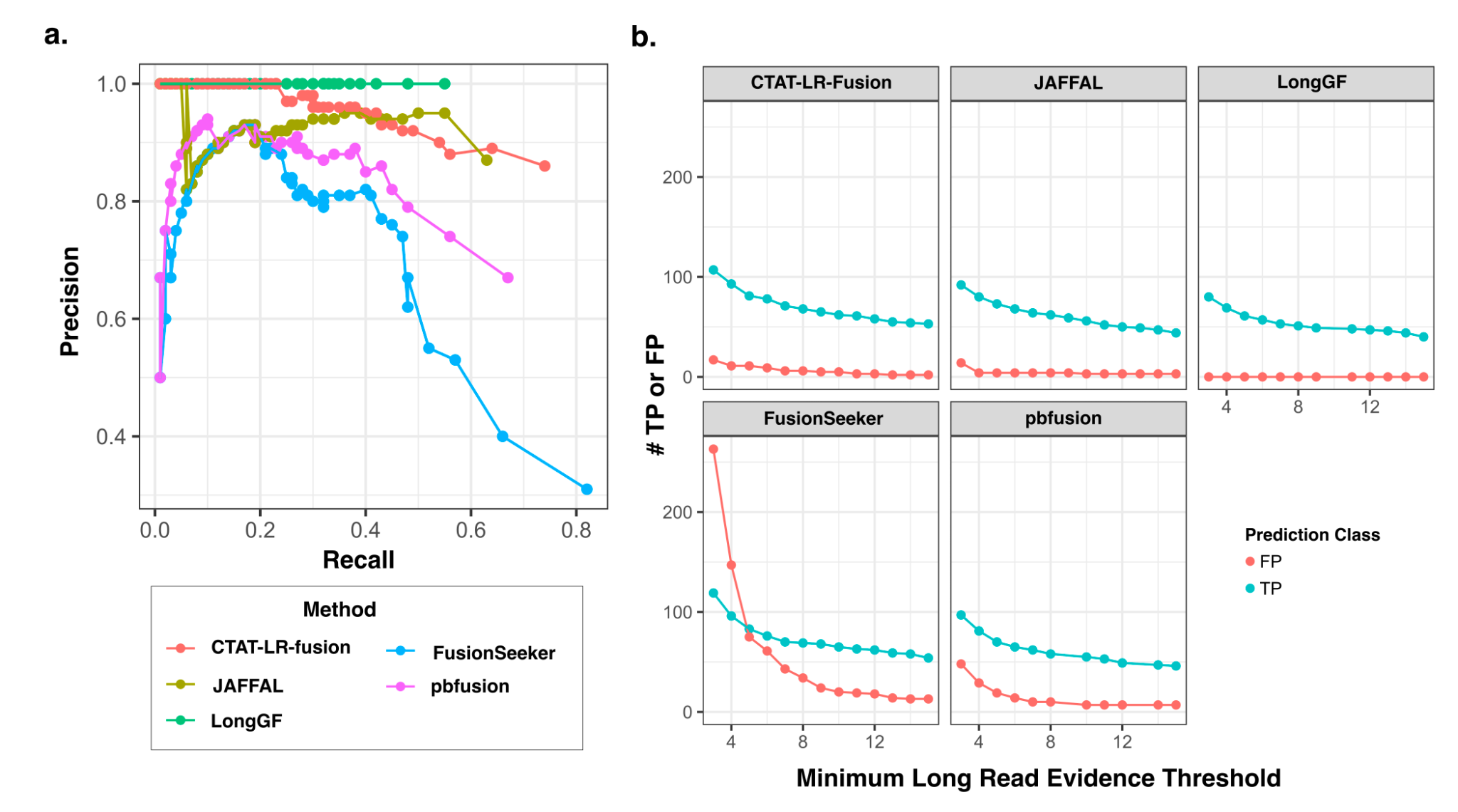

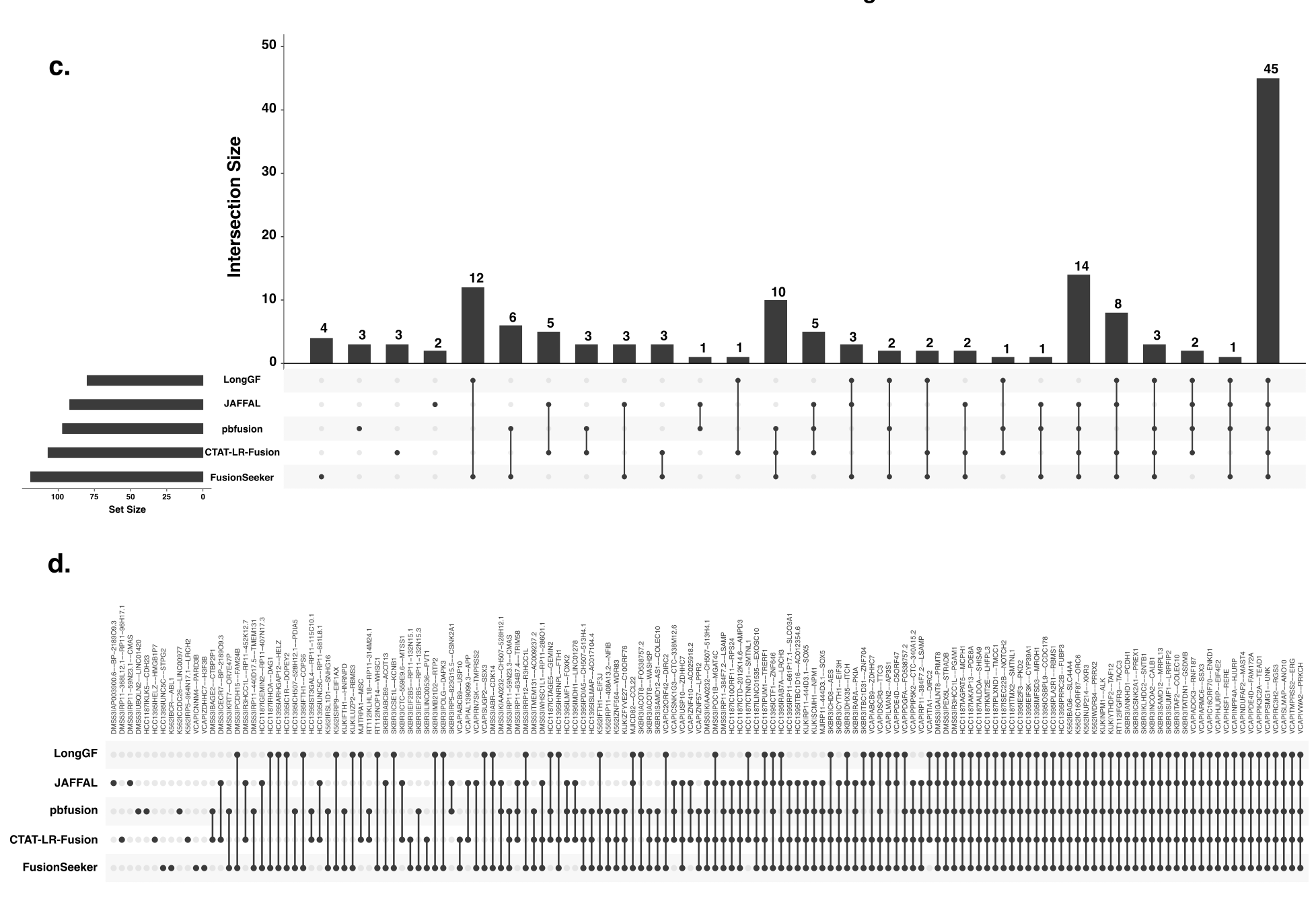

**Supplementary Figure 2**: Evaluation of fusions detected in nine DepMap cell lines. (a) Precision and recall measured at varied minimal long read evidence thresholds according to prediction method. (b) Counts of TP and FP according to minimum long read evidence count thresholds for each evaluated detection method. (c) Upset plot indicating counts of predictions found in common across combinations of prediction methods. (d) Fusions in the proxy truth set found according to the prediction method. Uniquely predicted fusions here are further supported by Illumina short reads.

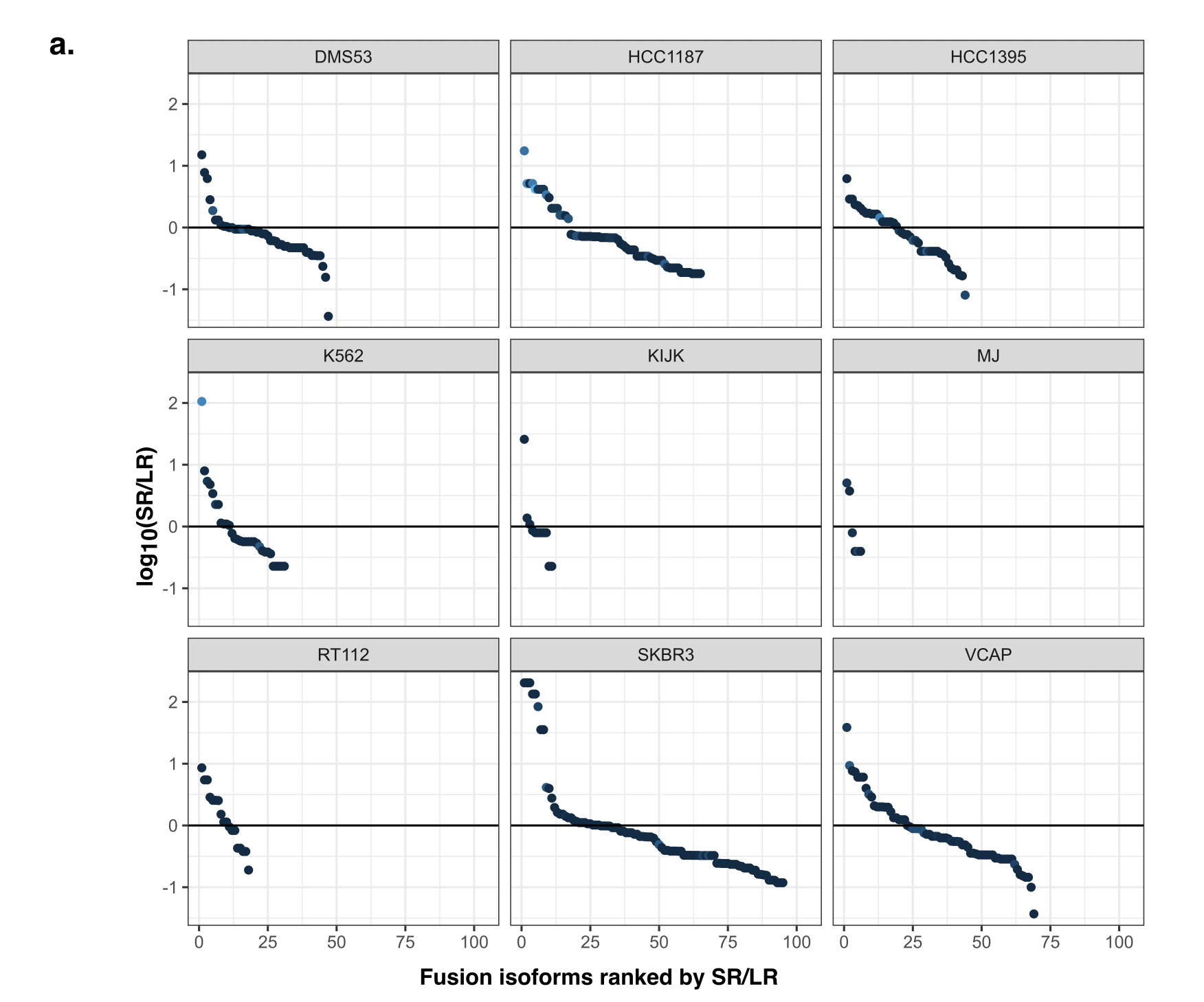

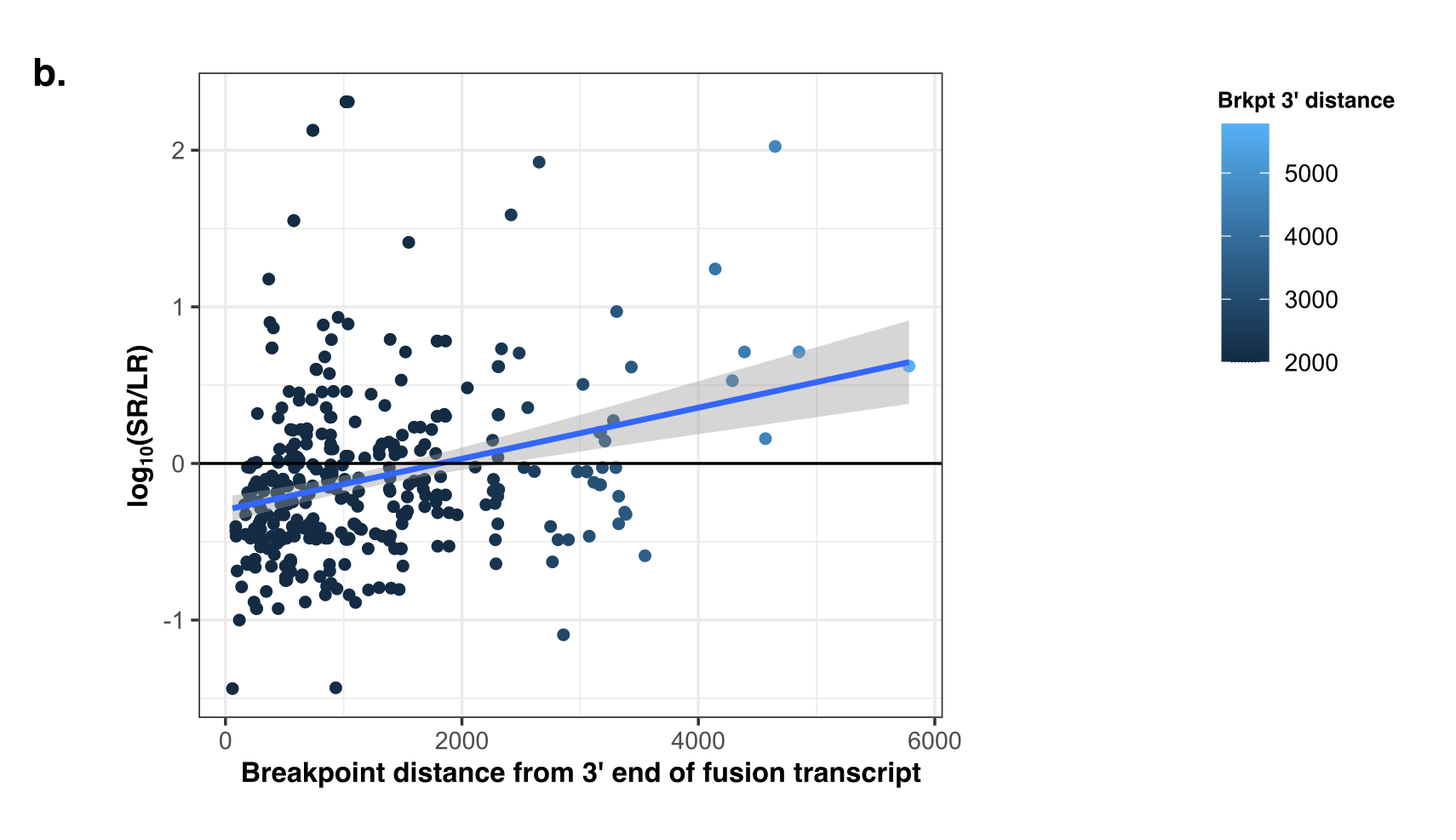

**Supplementary Figure 3: Short read (SR) vs. long read (LR) support for fusion evidence.** (a) Relative support of SR vs. LR support for fusion transcript isoforms, ranked by SR/LR and computed as log_10_(SR/LR) where SR and LR are normalized according to sequenced number of bases. (b) SR vs. LR support for fusion isoforms according to median distance of the breakpoint to the end of the read alignment.

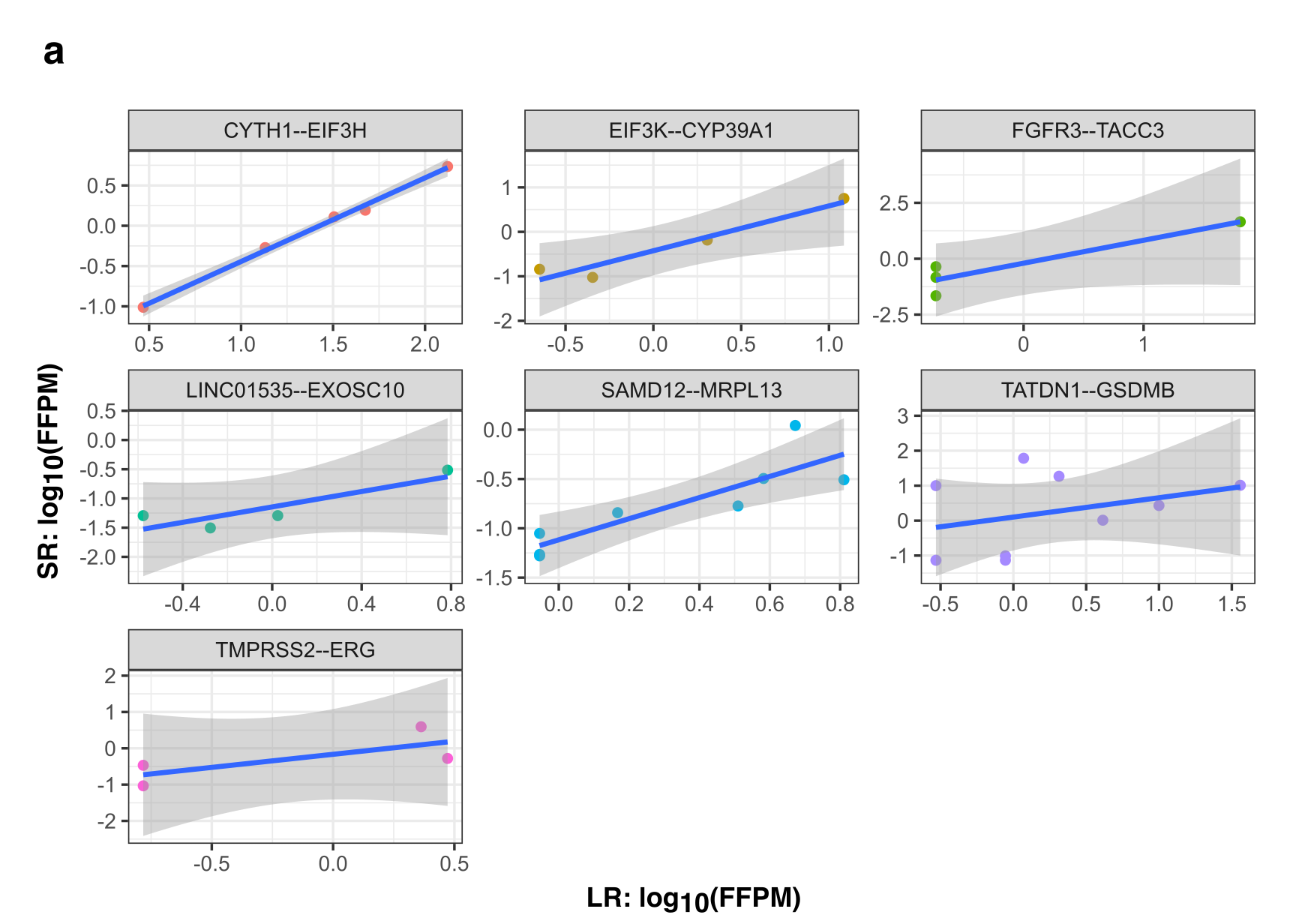

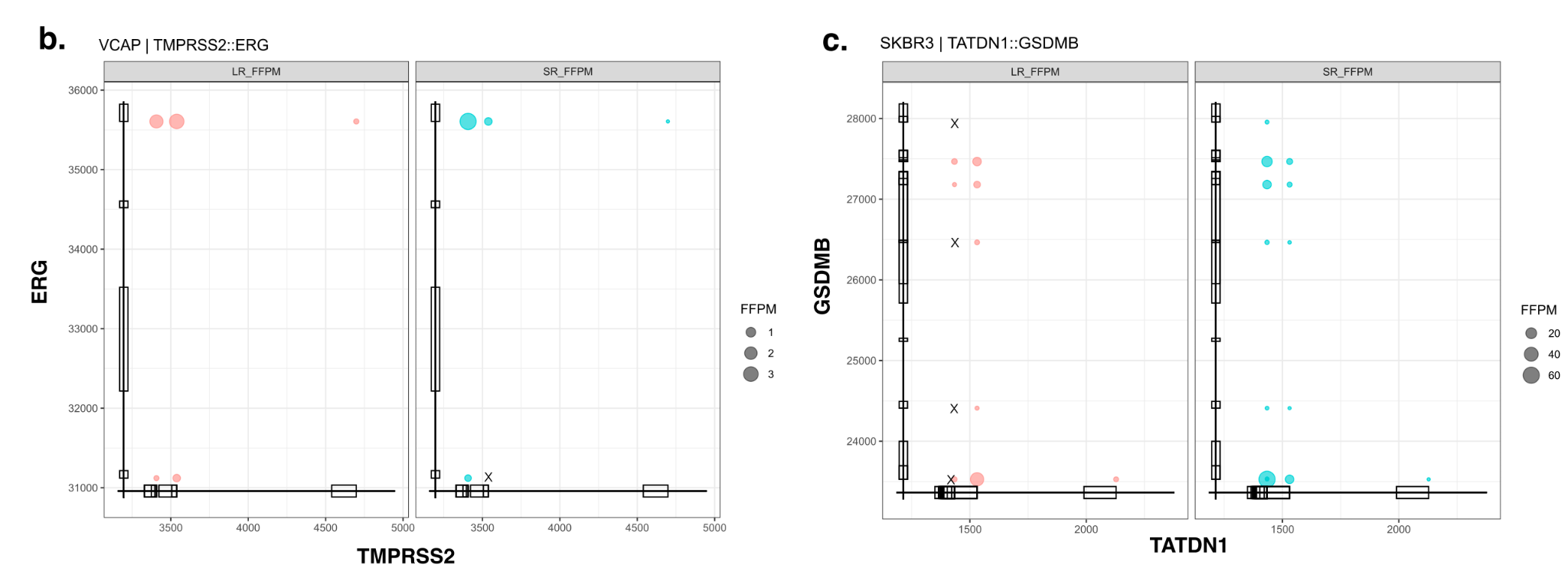

**Supplementary Figure 4: Comparison of short vs. long read support for fusion transcript isoforms.**  (a) Fusion expression evidence normalized as FFPM are compared for long reads (x-axis) and short reads (y-axis) for fusions CYTH1::EIF3H in SKBR3, LINC01535::EXOSC10 in HCC1187, SAMD12::MRPL13 in SKBR3, TATDN1::GSDMB in SKBR3, and TMPRSS2::ERG in VCaP. (b,c) Read alignment evidence quantified for fusion isoform breakpoints according to long (left, red) or short (right, blue) read RNA-seq: (b) TMPRSS2–ERG fusion in VCaP, (c) TATDN1–GSDMB fusion in SKBR3.
